# A tribal-scale phylogenomic perspective on ancestry and major lineages of the allotetraploid Hawaiian silversword alliance (Madieae, Compositae)

**DOI:** 10.64898/2026.07.26.740789

**Authors:** Bruce G. Baldwin, Susan Fawcett, Andrew Kostic, Bridget L. Wessa, William A. Freyman

**Affiliations:** Jepson Herbarium and Department of Integrative Biology, University of California, Berkeley, CA 94720-2465; National Tropical Botanical Garden, 3530 Papalina Road, Kalaheo, HI 96741; University and Jepson Herbaria, University of California, Berkeley, CA 94720-2465

**Keywords:** allopolyploidy, *Argyroxiphium*, Asteraceae, California flora, *Dubautia*, hybrid speciation, island biology, long-distance dispersal, Madiinae, *Wilkesia*

## Abstract

**Premise of the study:** Disentangling ancestry of insular plant lineages is important for resolving their evolutionary histories but often complicated by polyploidy, an overrepresented condition in remote island floras. A phylogenomic approach was used to study the origin of subgenomes of the tetraploid Hawaiian silversword alliance and advance understanding of this major example of adaptive radiation.

**Methods:** Sequences from 461 target-capture nuclear loci were phased to subgenomes of the tetraploid silversword alliance (Compositae) using assemblies from diploid North American tarweeds as clade references. Maximum-likelihood and multispecies-coalescent analyses of phased loci were conducted with and without representatives of diploid lineages throughout tribe Madieae.

**Key results:** An allopolyploid origin of the silversword alliance was corroborated, with Subgenome-A phased consistently to *Anisocarpus madioides* and Subgenome-B phased to diploid representatives of each of three other sublineages of the “Madia” lineage. Phylogenomic analyses indicate *Anisocarpus madioides* is more closely related to Subgenome-A than to other tarweeds, including *Anisocarpus scabridus*. Subgenome trees for the silversword alliance are highly congruent with each other and with structural genomic differences between lineages.

**Conclusions:** These results indicate an even closer relationship of the silversword alliance to extant California tarweeds than previously understood. *Anisocarpus madioides* has chromosomal and ecological traits anticipated in an ancestor of the silversword alliance, with the widest geographic range among perennial tarweeds and sticky bracts completely enveloping fruits. Within the silversword alliance, robust support for four major clades reinforces the phylogenetic value of chromosomal interchanges and taxonomic insights by Asa Gray regarding the enigmatic liana *Dubautia latifolia*.

---

Island plant lineages representing instructive examples of adaptive radiation are often ancestrally polyploid (Cerca et al., 2023). For example, hybridization and polyploidy have been documented in 40% of insular radiations globally in Compositae, the most diverse plant family on islands (Roeble et al., 2024). Polyploidy has been associated with higher success in long-distance dispersal to islands (Linder and Barker, 2014) and in subsequent insular diversification (Crawford et al., 2009). Achieving detailed evolutionary and biogeographic understanding in young, ancestrally polyploid radiations has been challenging, however, in part because of lack of access to low-copy nuclear genes in polyploids without overcoming the difficulty of assigning loci to subgenomes (see Rothfels, 2021). This limitation may be particularly acute for resolving ancestry of island radiations if extra-insular relatives stemming from the same polyploidization event are extinct or unknown.

The Hawaiian silversword alliance (Madiinae--Madieae, Compositae) is a prominent example of adaptive radiation (Figures 1 and 2; Schluter, 2000), with experimental and molecular phylogenetic evidence indicating an allotetraploid ancestry (Barrier et al., 1999). Specifically, floral homeotic gene copies cloned from each subgenome of the silversword alliance were resolved by Barrier et al. (1999) as more closely related to different diploid sublineages of the continental “Madia” lineage (Figures 3 and 4; Baldwin, 1996, 2003) than to each other, in support of a hybrid origin of the polyploid condition in silverswords. A subsequent phylogenetic analysis of cloned growth regulator genes corroborated an allopolyploid ancestry of the silversword alliance involving the “Madia” lineage, with somewhat different resolution of relationships of subgenomic copies to the continental taxa (Remington and Purugganan, 2002). Although these and earlier studies of the tarweed--silversword relationship have indicated an ancestry of the silversword alliance from within the “Madia” lineage of North American tarweeds (e.g., Baldwin et al., 1991; Carr et al., 1996; Baldwin and Wessa, 2000; Baldwin, 2003a), precise relationships of the silversword alliance subgenomes within the “Madia” lineage have remained uncertain. Similarly, considerable resolution of the silversword alliance radiation has been achieved with cytogenetic, nuclear ribosomal DNA (nrDNA), plastid, and nuclear regulatory gene data (e.g., Carr and Kyhos, 1986; Barrier et al., 1999; Carr, 2003; Baldwin et al., 2021), but extensive cytonuclear discordance, other phylogenetic incongruence, and lack of robust support for some relationships have made evident the need for a phylogenomic perspective involving a large number of nuclear genes to achieve a more refined understanding of silversword evolution.

**Fig. 1.**
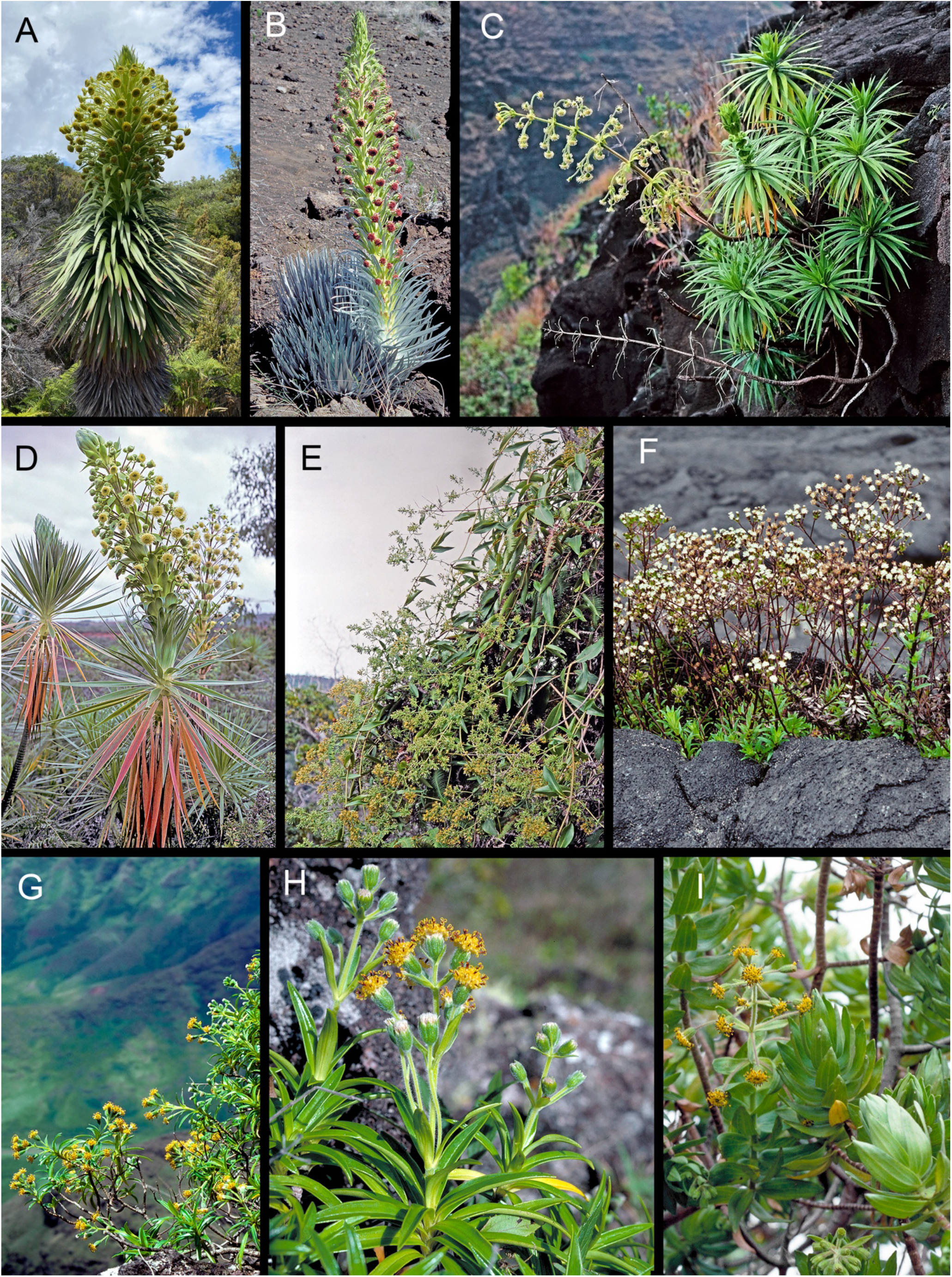
Representatives of the endemic Hawaiian silversword alliance (Argyroxiphium, Dubautia, Wilkesia; Madiinae). (A) Argyroxiphium grayanum. (B) A. sandwicense subsp. sandwicense. (C) Wilkesia hobdyi. (D) W. gymnoxiphium. (E) Dubautia latifolia. (F) D. scabra subsp. scabra. (G, H) D. herbstobatae. (I) D. arborea. Photo credits: (A) Zach Pezzillo. (B--I) Gerald D. Carr.

**Fig. 2.**
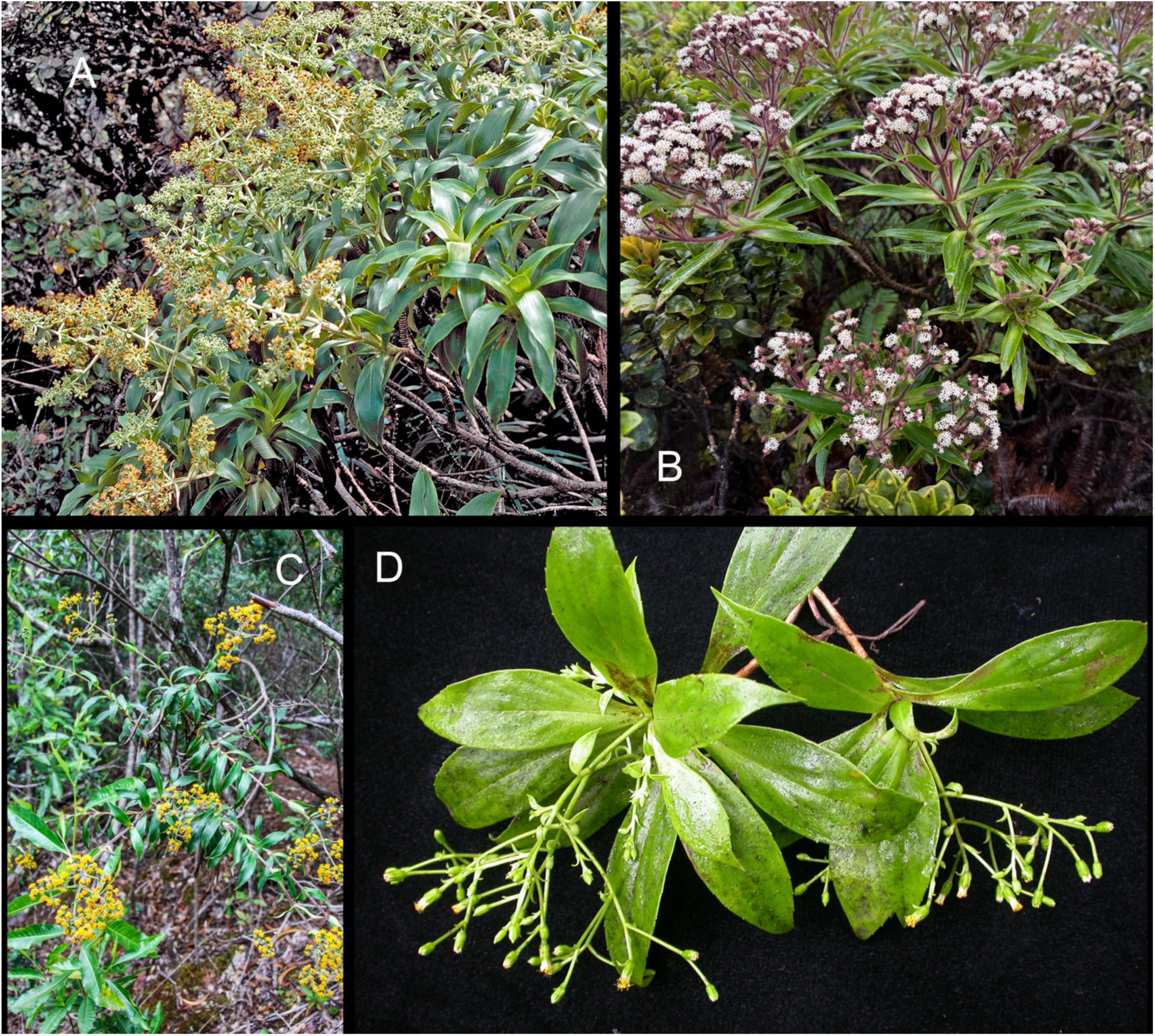
Additional representatives of the Hawaiian silversword alliance (Madiinae), in *Dubautia* sect. *Dubautia*. (A) *Dubautia plantaginea* subsp. *plantaginea*. (B) *D. paleata*. (C) *D. laevigata*. (D) *D. knudsenii* subsp. *filiformis*. Photo credits: (A) Gerald D. Carr. (B) C. Newfield. (C, D) Kenneth R. Wood.

**Fig. 3.**
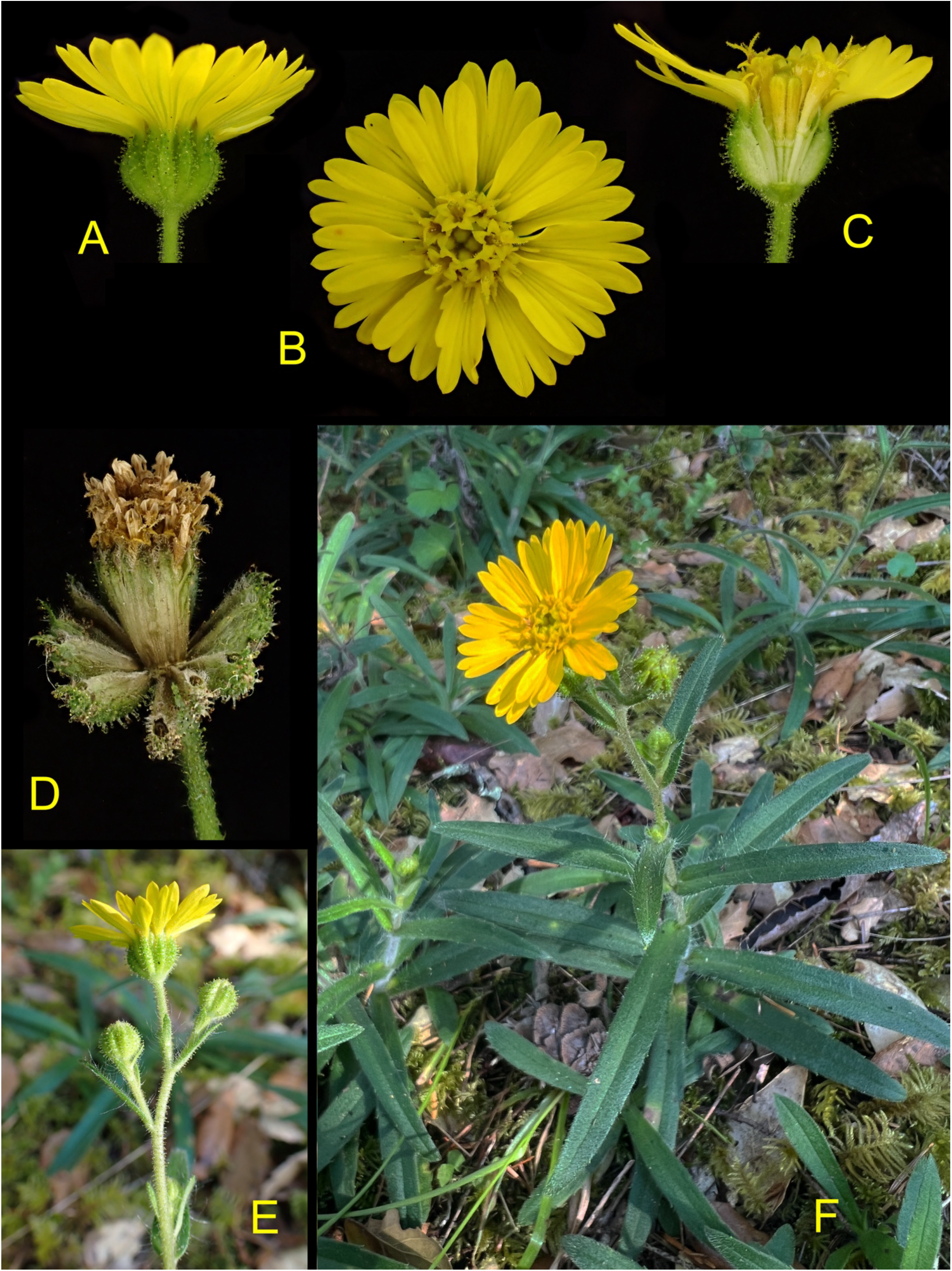
*Anisocarpus madioides*, a perennial North American tarweed in the “Madia” lineage (see Figure 8) of Madiinae. (A) Flowering head, lateral view. (B) Flowering head, frontal view. (C) Flower head longisection, showing disk florets and glandular involucral bracts (phyllaries), each enveloping the base of a ray floret. (D) Fruiting head, with reflexed ray fruits, each tightly enveloped by a sticky phyllary, and post-anthesis pistillate-sterile disk florets collectively surrounded by a ring of connate receptacular bracts. (E) Young capitulescence, lateral view. (F) Plant habit, in forest understory. Photo credit: (A--F) Susan Fawcett.

**Fig. 4.**
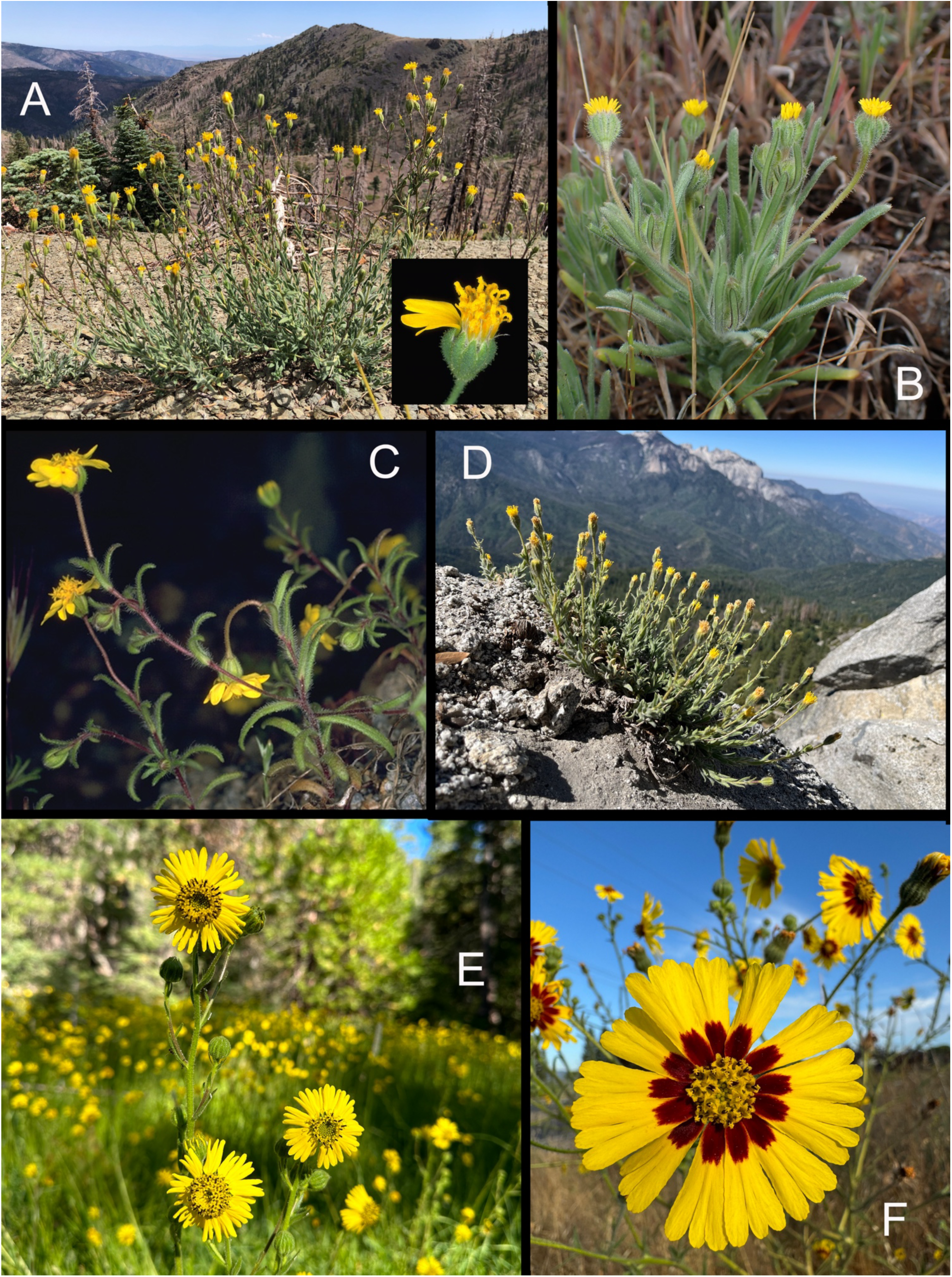
North American tarweed representatives of major sublineages of the “Madia” lineage (Madiinae), in addition to *Anisocarpus madioides* (Figure 3). (A) *Anisocarpus scabridus* [= *Raillardiopsis scabrida*], habit and flowering head inset. (B) *Jensia yosemitana*. (C) *Harmonia nutans*. (D) *Carlquistia muirii*. (E) *Kyhosia bolanderi*. (F) *Madia elegans*. Photo credits: (A) habit – Jen Pagel; head inset – Gerald D. Carr. (B) Preston Ernest. (C) Bruce G Baldwin (D) Dean Lyons. (E) Matthew Berger. (F) Jacob (earthbound_adverturer).

Here, a phylogenomic approach to resolving ancestry of the tetraploid silversword alliance was pursued using target enrichment of hundreds of nuclear loci and phasing of sequences from these loci to each of the two silversword subgenomes using diploid tarweed species of the “Madia” lineage as clade references, followed by phylogenomic analysis of these phased sequences with representatives of genera throughout the tarweed tribe, Madieae. We address the following questions: (1) Do extant North American tarweeds serve as effective clade references for phasing of sequence reads to silversword alliance subgenomes, or are the subgenomes insufficiently divergent from one another or too dissimilar to genomes of modern tarweeds, after 5+ million years since common ancestry (Landis et al., 2018)? (2) Does any evidence of subgenome relationships to North American tarweeds provide potential insight into ecological characteristics of the colonizing ancestor of the silversword alliance, or are relationships to extant tarweeds too distant to draw any conclusions? (3) Is any resolution of the silversword alliance radiation achieved from phased sequences and, if so, are results congruent across subgenomes or pertinent to any outstanding evolutionary questions? (4) At a broader-scale, are relationships across Madieae resolved and, if so, what can be inferred about the deeper evolutionary and ecological history of the tribe?

## MATERIALS AND METHODS

### Taxon sampling

Sampling of the Hawaiian silversword alliance included representative species of all genera (*Argyroxiphium* DC., *Dubautia* Gaudich., *Wilkesia* A.Gray) and taxonomic sections (Carr, 1985), and all known structural genomes (Carr and Kyhos, 1986; Carr, 2003). For the remaining diversity in the primarily western North American tribe Madieae, all recognized subtribes and all genera with any diploid diversity (i.e., all but one genus, *Hemizonella* (A.Gray) A.Gray) were included, with attention to sampling across the root node of genera, based on prior data of relationships from more detailed sampling (Baldwin, 2003a, unpublished data; the authors, in prep.). Outside the silversword alliance, all sampled taxa were judged to be diploid (of paleopolyploid ancestry, with polyploidization pre-dating tribe Madieae; Barker et al., 2016b; Huang et al., 2016; Zhang et al., 2021), based on high congruence between reconstructions of chromosome evolution (Baldwin and Wessa, 2000; Carr, 2003) and distribution of long-paralog warnings in HybPiper (see below). For continental tarweeds of the “Madia” lineage, which includes the closest relatives of the silversword alliance based on all previous studies, all recognized diploid species were included, following analyses by the authors, to be published elsewhere, indicating that polyploid continental taxa in the “Madia” lineage represent distinct polyploidization events from the polyploidization in the ancestral lineage of the silversword alliance. For subgenomic phasing, diploid reference taxa represented each of the six robust sublineages of the “Madia” lineage, based on preliminary phylogenomic analyses of the target enrichment data. Voucher information for sampled taxa is in Appendix 1.

### Target-capture of nuclear loci

Genomic DNAs of sampled taxa were extracted from young leaves as described by Baldwin et al. (2021), using organellar isolation, a cetyltrimethylammonium bromide (CTAB) protocol modified from Doyle and Doyle (1987), or the DNeasy Plant Mini Kit (QIAGEN, Germantown, MD). For summer-flowering California tarweeds (in Madiinae), leaves for DNA extraction were harvested from basal rosettes of young greenhouse-grown plants to avoid polysaccharides and glandular exudates of mature foliage. Genomic DNAs were purified on Monarch silica columns (New England BioLabs, Ipswich, MA). Library preparation, target-capture, and high-throughput sequencing were performed by Arbor Biosciences (Ann Arbor, MI). DNAs were sheared to 400-600 nt average insert length by sonication and dual-indexed using TruSeq adapters (Illumina Inc., San Diego, CA). Resulting libraries were target enriched (8--12 libraries/reaction) and pooled for Illumina NovaSeq S4 PE150 (paired-end 150 bp) sequencing, with tetraploid (silversword alliance) samples sequenced to twice the depth of coverage as diploids (1--2 Gbp/library). Baits for target capture (19,854 bait sequences, 80 nt each, spaced about every 28 nt for 3× tiling) for 461 putative single-copy nuclear loci (> 900 bp each; ∼600 kb total) were designed to develop a custom MYbaits probe library (Arbor Biosciences) in this study from leaf transcriptomes of *Dubautia latifolia* and *Wilkesia gymnoxiphium* (M. S. Barker, R. H. Robichaux, B. G. Baldwin, unpublished data) using MarkerMiner 1.0 (Chamala et al., 2015).

### Gene assembly

Sequencing reads were quality-filtered and trimmed to remove adapters and polyG tails using fastp (Chen et al., 2018) prior to gene assembly. For diploid samples, cleaned sequencing reads were target-mapped to gene directory using BWA (Li and Durbin, 2009) and assembled into contigs using SPAdes (Prjibelski et al., 2020), within the HybPiper 2 pipeline (Johnson et al., 2016; https://github.com/mossmatters/HybPiper). Contig stitching, where necessary, was done in reference to target protein sequences using Exonerate (in HybPiper), with retrieval of both exons and supercontigs (exons + flanking regions) using the intronerate.py script. For tetraploid (silversword alliance) samples, cleaned sequencing reads were mapped to target references using samtools (Danecek et al., 2021) and matched to gene assemblies of diploid species representing the six major sublineages of the “Madia” lineage (i.e., *Anisocarpus madioides*, *A. scabridus*, *Carlquistia muirii* [*Baldwin 618* (DAV)] (representing the sublineage that also includes *Madia*)*, Harmonia guggolziorum*, *Jensia yosemitana*, and *Kyhosia bolanderi*) using BBSplit in HybPhaser (Nauheimer et al., 2021). The resulting clade association summary table, with percentage of unambiguously assigned reads to each diploid species, helped to guide subsequent selection of references for phasing of silversword alliance reads to subgenome (Figure 5). Four pairs of clade references were chosen from the “Madia” lineage of North American Madiinae to represent each of four sublineages (and all perennial continental taxa) therein for subsequent read phasing: *Anisocarpus madioides* and *A. scabridus*, *A. madioides* and *Carlquistia muirii*, *A. madioides* and *Kyhosia bolanderi*, and *Carlquistia muirii* and *Kyhosia bolanderi*. Sequencing reads of the silversword alliance phased to each of the four pairs of clade references were then assembled using HybPiper and HybPhaser.

**Fig. 5.**
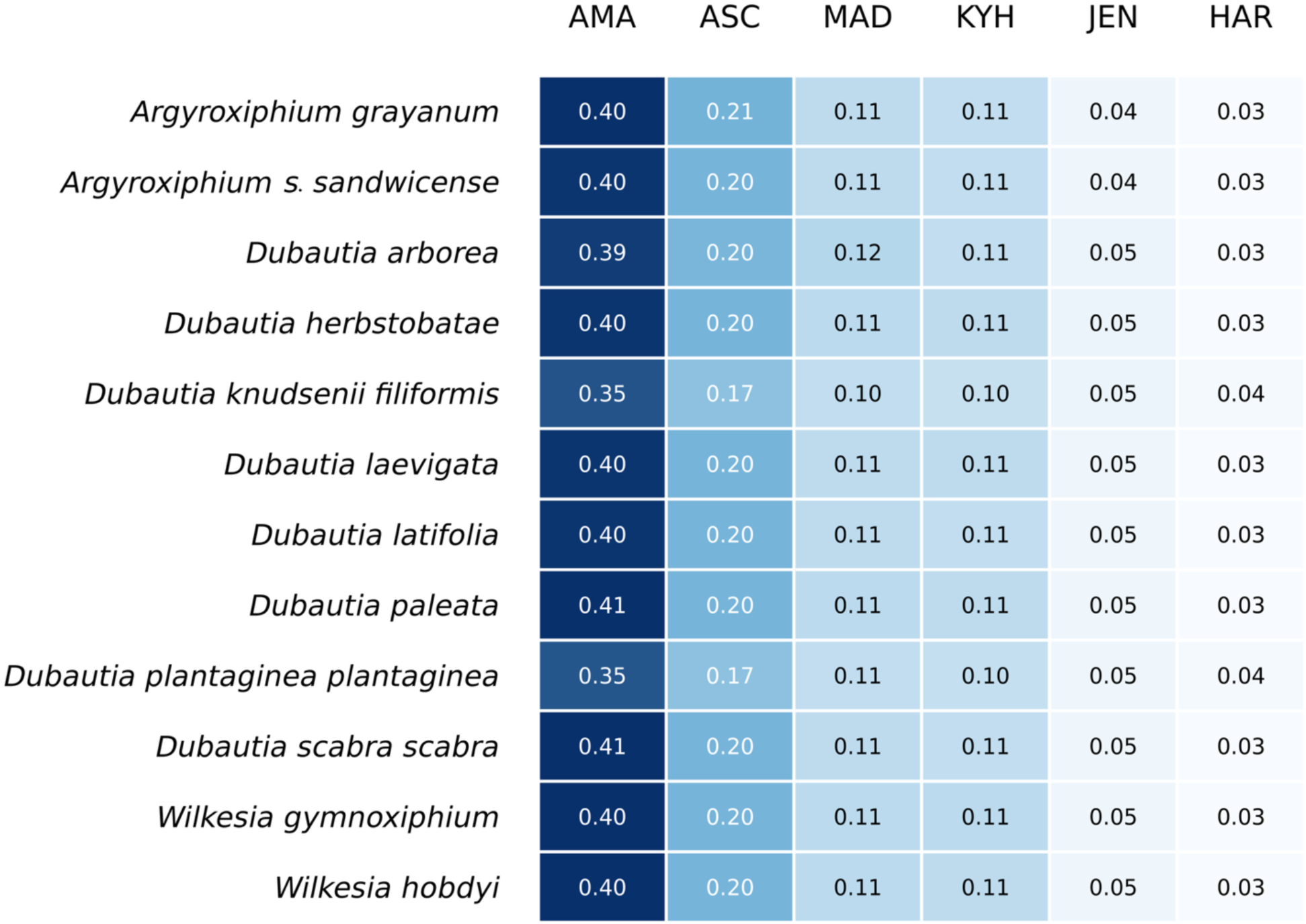
Heatmap showing proportion of sequencing reads of the silversword alliance mapped to reference clades. Proportions are shown for association of reads from each of the 12 sampled Hawaiian taxa mapped to each of six diploid tarweed clade references: *Anisocarpus madioides* (AMA), *A. scabridus* (ASC), *Carlquistia muirii* (MAD)*, Kyhosia bolanderi* (KYH), *Jensia yosemitana* (JEN), and *Harmonia guggolziorum* (HAR).

### Phylogenomic analyses

Gene assemblies for each target region were aligned using MAFFT v7.525 (Katoh and Standley, 2013), for exons only and for supercontigs. Assemblies of the silversword alliance resulting from phasing of reads to each pair of clade references were aligned together for unrooted phylogenomic analyses. In addition, assemblies from reads of the silversword alliance phased to one of the diploid reference species in each pair were merged with assemblies of diploid samples representing genera across tribe Madieae for rooted phylogenomic analyses. Problematically aligned positions and loci (e.g., with excessive gaps) were removed prior to phylogenomic analysis using trimAl (Capella-Gutiérrez et al., 2009). Maximum-likelihood (ML) analysis of concatenated genes was conducted using IQ-TREE v3 (Nguyen et al., 2015), with the best-fit model of sequence evolution estimated by ModelFinder (Kalyaanamoorthy et al., 2017) and ultrafast bootstrap support of clades approximated by UFBoot (Minh et al., 2013). IQ-TREE results for each gene were used in subsequent multispecies-coalescent (“species tree”) analysis using ASTRAL v5.7.8 (Zhang et al., 2018). The sole member of Venegasiinae (*Venegasia carpesioides*) was used to root the trees based on results of Zhang et al. (2021).

### Trait mapping

Morphological character evolution across tribe Madieae was mapped onto ultrametric trees with the multispecies-coalescent (ASTRAL) topologies using maximum-likelihood in phytools (Revell, 2024). Trees used for trait mapping represented the main ASTRAL topologies for supercontig assemblies of reads phased to each pair of clade references. Maximum-likelihood branch lengths for the concatenated 461-locus nuclear dataset were optimized for the ASTRAL tree topologies using IQ-TREE v3 and then an ultrametric tree was fitted to the ML tree using penalized likelihood (Sanderson, 2002) in chronos (Paradis, 2013), within the ape (Analyses of Phylogenetics and Evolution) package in R (Paradis and Schliep, 2019). Characters examined included those of potential ecological importance, for example, in dispersal and colonization. Morphological traits were assessed directly from specimens, and are well-documented in protologues, monographs, and floras (e.g., Carr 1985; Baldwin et al. 2012). Self-incompatibility or lack thereof was scored based on experimental crosses by the first author or others (Keck 1959; Ornduff 1966; Mooring 1975; Wilken 1975; Carr et al. 1986; Kyhos et al. 1990). Chromosome-number evolution in tribe Madieae was also estimated for the “Madia” lineage, pruned from the same ultrametric trees, using chomEvol 2.0 (Glick and Mayrose, 2014), with selection of the best fitting model for analysis (CONST_RATE: constant rate of chromosome gains and losses and genomic duplications for all trees) based on log-likelihood and AIC scores, and with *Holozonia* Greene as outgroup.

### Computational resources

Analyses were conducted on Savio, the high performance computing cluster at the University of California, Berkeley. Claude (Anthropic Sonnet 4.6 2026; www.anthropic.com) was used for troubleshooting code, basic scripting, and organizing and tracking pipeline status on the Savio cluster. Scripts and additional methodological details available on GitHub: github.com/susanfawcett/Madieae/tree/Allopolyploid_Origin_Silverswords

## RESULTS

### Diploid gene assembly

DNA sequences were recovered for 459 to 461 of the 461 target genes across all but two sampled Madieae taxa (456 genes for *Centromadia pungens* and 457 for *Lasthenia californica*), with > 75% of reference target sequence recovered for each of 458 to 461 genes in > 75% of samples, and for each of 450 to 461 genes in > 90% of samples, i.e., all but *Blepharipappus scaber* (440), *Lasthenia californica* (419), *Lasthenia glaberrima* (445), and *Syntrichopappus fremontii* (447). In all, 583 to 603.9 kb (*x̅* = 596.1 kb) of target sequence were obtained per sample, except for *Blepharipappus scaber* (572 kb) and *Lasthenia californica* (542.4 kb). Depth of coverage for target sequences ranged from ∼ 134 to 5642× (*x̅* = 1502×) except for *Blepharipappus scaber* (∼93×). For the four clade references used for phasing sequence data of silversword alliance taxa, 460 or 461 target genes were sequenced, with > 75% of target sequence recovered in >= 459 genes (599.3 to 599.9 kb total, at ∼ 372 to 2229× (*x̅* = 1815×) depth of coverage).

### Polyploid gene assembly

Results of clade association of silversword alliance sequencing reads to assemblies of diploid references for each of the six main sublineages of the “Madia” lineage (Figure 5) indicated the following mean level of unambiguous association of sequencing reads to each of the following reference taxa (in descending order): *Anisocarpus madioides* (0.392), *A. scabridus* (0.199), *Carlquistia muirii* (0.111), *Kyhosia bolanderi* (0.107), *Jensia yosemitana* (0.047), and *Harmonia guggolziorum* (0.034). *Anisocarpus madioides*, with approximately twice as many unambiguously assigned reads as the diploid reference with the second highest level of unambiguously assigned reads (*A. scabridus*), was subsequently paired with *A. scabridus*, *C. muirii*, and *K. bolanderi* to explore performance of these clade representatives for sequencing-read phasing. The two diploid references outside *Anisocarpus* Nutt. (i.e., *C. muirii* and *K. bolanderi*) were also paired to assess their utility for sequencing-read phasing. Phasing statistics for each of these four pairs of clade references (Figure 6) showed again that *A. madioides* had the highest level of unambiguously assigned reads, with *A. scabridus* having a greater number of unambiguously assigned reads (ca. 2/3 that of *A. madioides*) than either *C. muirii or K. bolanderi* (each < 1/2 that of *A. madioides*) when paired with *A. madioides*. After sequence alignment and alignment trimming, all 461 loci were kept for silversword alliance exon assemblies and for silversword alliance supercontig assemblies from phasing to *A. madioides* and *A. scabridus*, and 48--58% of loci were kept for supercontig assemblies phased to *A. madioides* and *K. bolanderi* (267 loci), *A. madioides* and *C. muirii* (264 loci), and *C. muirii* and *K. bolanderi* (223 loci).

**Fig. 6.**
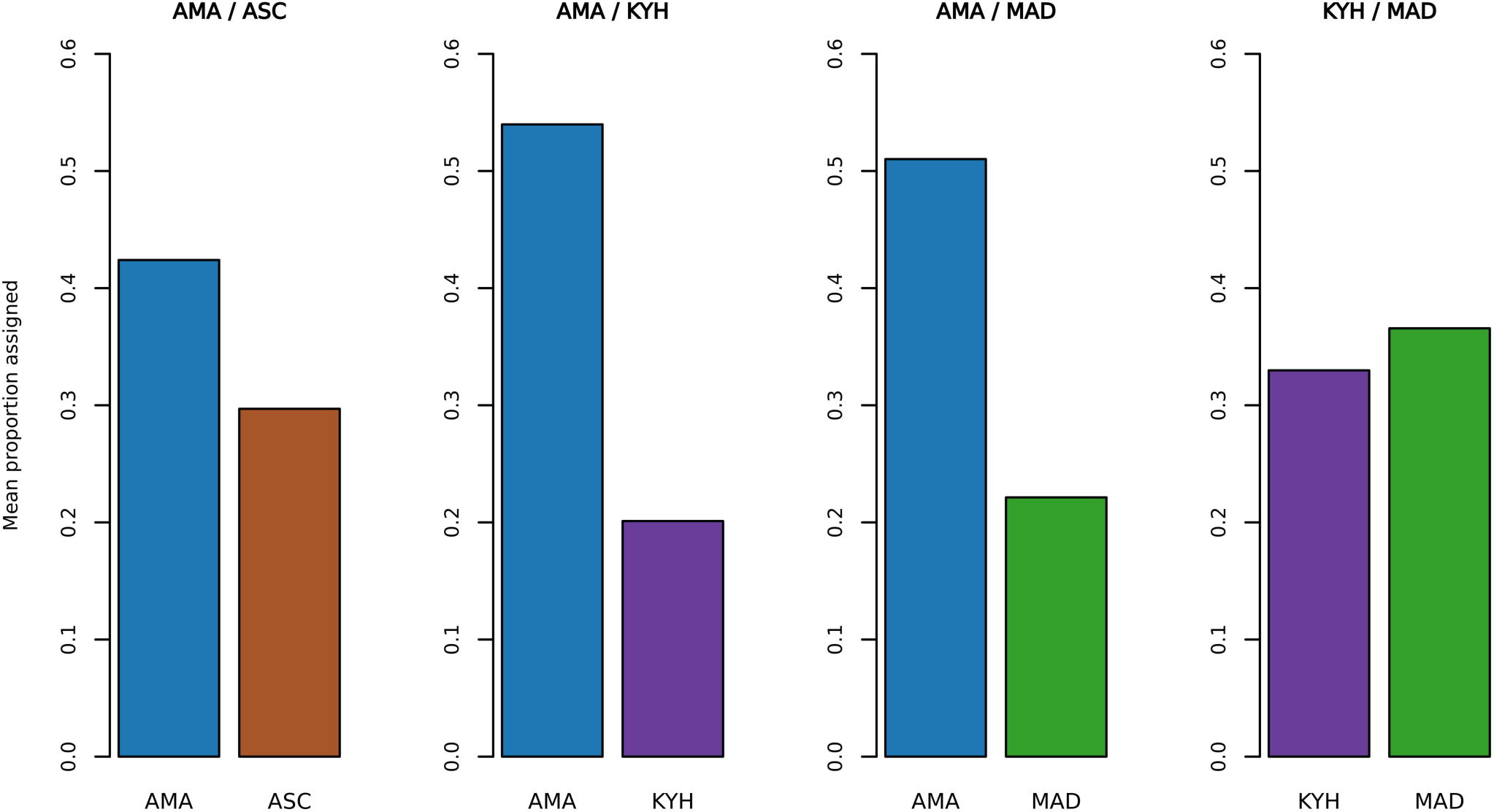
Clade association of silversword alliance sequencing reads to four pairs of diploid tarweed clade references (left to right): *Anisocarpus madioides* and *A. scabridus* (AMA / ASC); *A. madioides* and *K. bolanderi* (AMA / KYH); *A. madioides* and *Carlquistia muirii* (AMA / MAD); and *K. bolanderi* and *Carlquistia muirii* (KYH / MAD).

### Subgenome-level phylogenomic resolution in the silversword alliance

Phylogenomic analyses of the silversword alliance that included supercontig assemblies phased to both *Anisocarpus madioides* and one of the other three diploid species of the “Madia” lineage paired with it (*A. scabridus*, *Carlquistia muirii*, or *Kyhosia bolanderi*) resulted in unrooted ASTRAL trees with identical or nearly identical, fully resolved silversword alliance clades separated by an internal branch, with the two clades representing gene assemblies phased to different diploid references (Figure 7A--C). In contrast, phylogenomic analyses of assemblies phased to the paired references *C. muirii* and *K. bolanderi* (without *A. madioides*) resulted in unrooted ML and ASTRAL trees with assemblies of the same silversword alliance species (sample) phased to different diploid references consistently being resolved together (Figure 7D), as expected with predominant phasing of both sets of assemblies to the same subgenome. ASTRAL trees for each of the four sets of phased assemblies were completely congruent with one another but differed substantially from ML trees (see Supplementary Information).

**Fig. 7.**
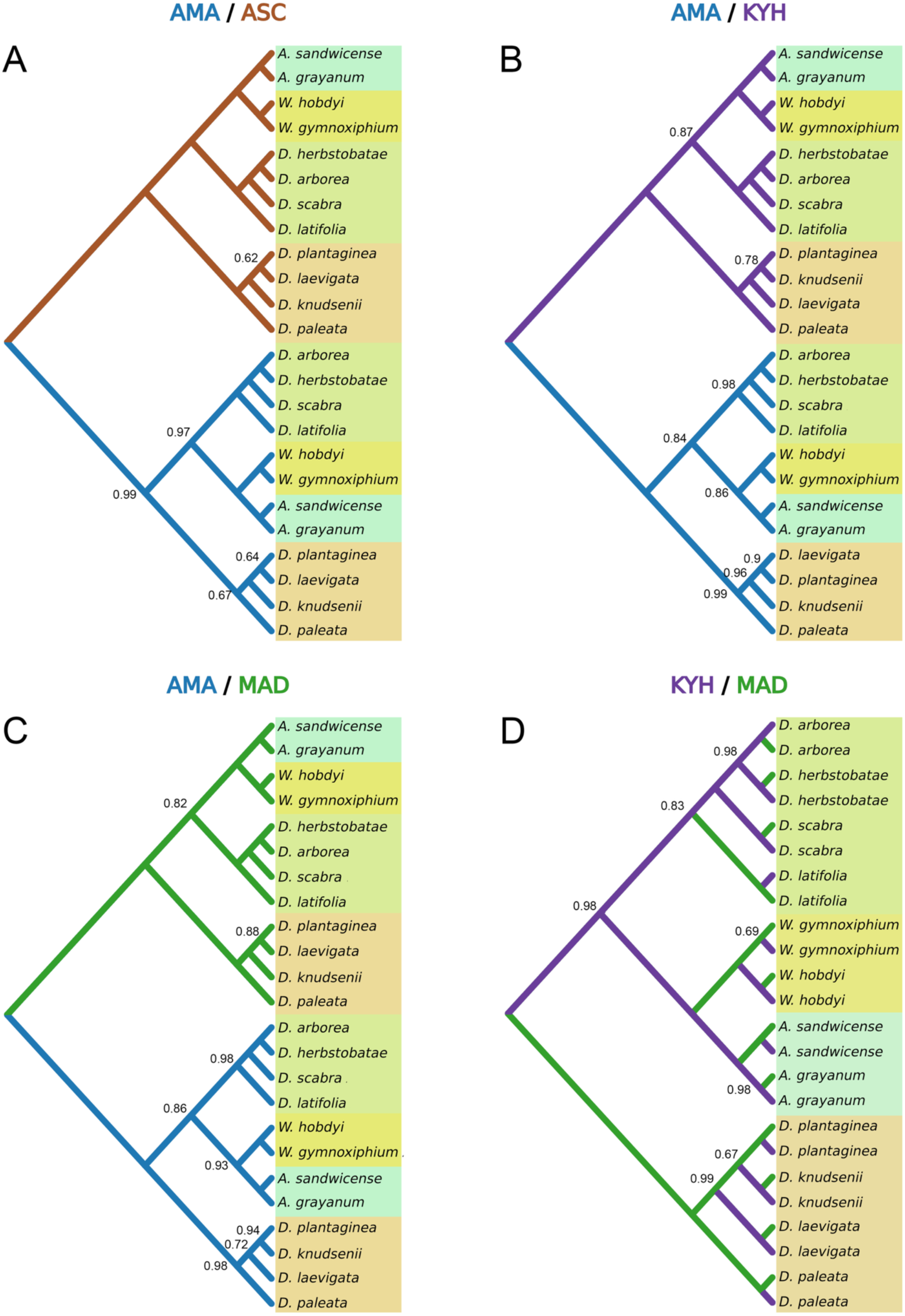
Unrooted multispecies coalescent (ASTRAL) trees of representatives of all genera and known chromosome arrangements in the Hawaiian silversword alliance for nuclear supercontig assemblies from sequencing reads phased to each of the following four pairs of tarweed references in Figure 6. (A) *Anisocarpus madioides* and *A. scabridus* (AMA / ASC). (B) *A. madioides* and *K. bolanderi* (AMA / KYH). (C) *A. madioides* and *Carlquistia muirii* (AMA / MAD). (D) *K. bolanderi* and *Carlquistia muirii* (KYH / MAD). Local posterior probabilities are shown at tree nodes if < 1.0. Colored blocks of taxa correspond to the following consistently resolved groups: *Argyroxiphium* (blue-green), *Dubautia* sect. *Dubautia* (tan), *Dubautia latifolia* + *D.* sect. *Railliardia* (olive-green), and *Wilkesia* (yellow-green). Note that branch color corresponds to color of abbreviated name (above tree) for reference taxon to which silversword alliance sequence reads were phased.

### Tribal-scale phylogenomic results

Phylogenomic analyses of individual phased sets of silversword alliance assemblies together with diploid representatives of genera across tribe Madieae yielded highly similar multispecies-coalescent trees, with the silversword alliance resolved as monophyletic and nested within a well-supported “*Madia*” lineage. For analyses with silversword alliance exon or supercontig assemblies phased to *Anisocarpus madioides*, the silversword alliance was always placed strongly sister to *A. madioides* in multispecies-coalescent trees (Figure 8). Phylogenomic analyses of exons or supercontigs assembled from Hawaiian reads phased to any of the three species *Anisocarpus scabridus*, *Carlquistia muirii*, or *Kyhosia bolanderi* paired in clade associations with *A. madioides* consistently and robustly placed the silversword alliance with *A. madioides* plus a strongly supported clade of *Carlquistia* B.G.Baldwin, *Harmonia* B.G.Baldwin, *Jensia* B.G.Baldwin, and *Madia* Molina. Local posterior probability (LPP) for a clade uniting *A. madioides* with *Carlquistia*, *Harmonia*, *Jensia*, and *Madia* was inconclusive (0.38--0.82 LPP), with three of six analyses instead resolving *A. madioides* as sister to the Hawaiian taxa (1.0 LPP) (Figure 9). Among diploids, high resolution and support for relationships throughout tribe Madieae was obtained, with few weakly supported lineages and most of these resolved with stronger support in analyses based on a more restricted (exon) or a more extensive (supercontig) locus assembly.

**Fig. 8.**
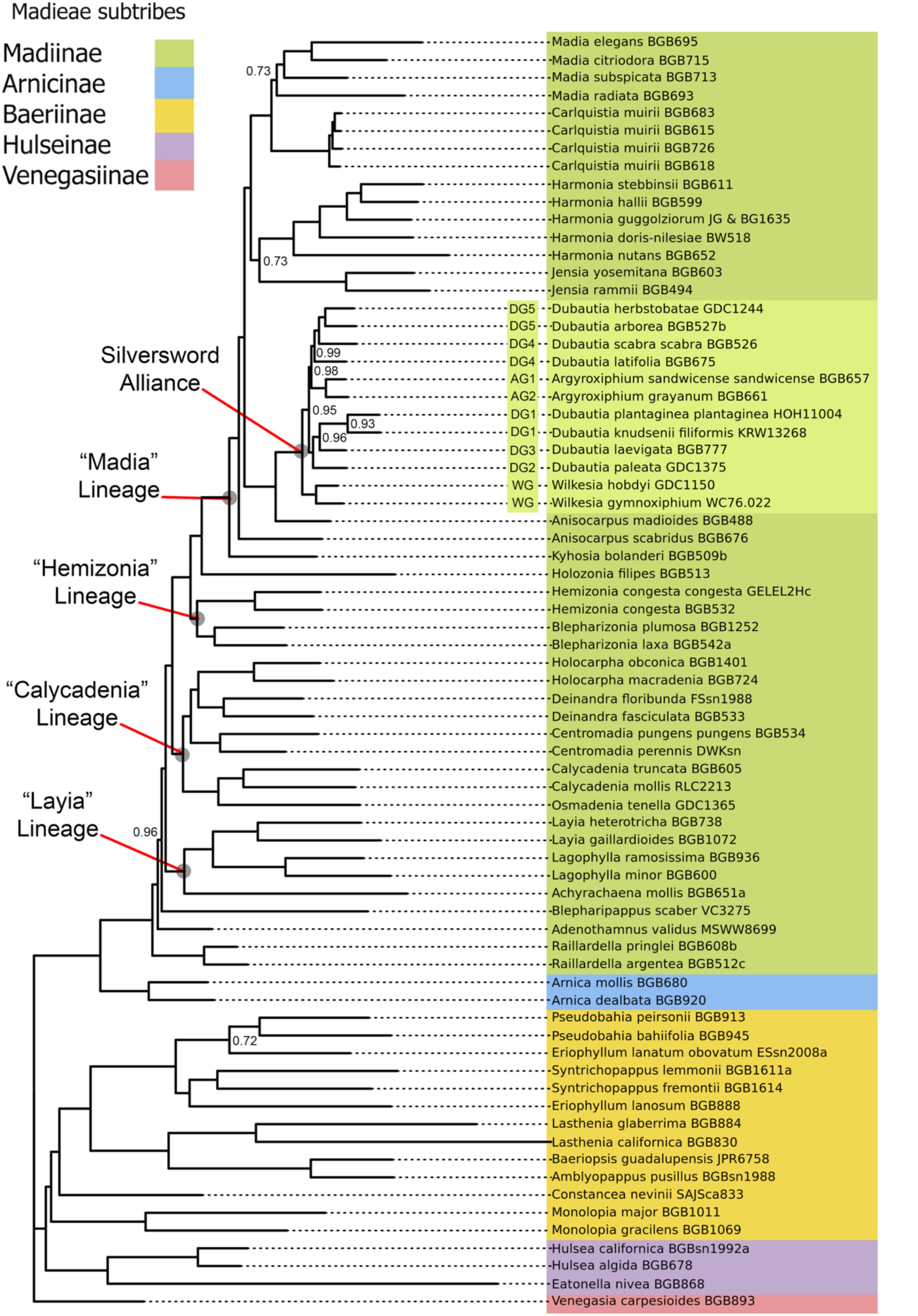
Maximum-likelihood phylogram of the multispecies coalescent (ASTRAL) tree topology for nuclear supercontig assemblies of representatives of all genera with diploid species in tribe Madieae plus representatives of all genera and known chromosome arrangements in the Hawaiian silversword alliance. Gene assemblies of the Hawaiian silversword alliance are from reads phased to *Anisocarpus madioides*, when paired with *A. scabridus*. Local posterior probabilities are shown at tree nodes if < 1.0. Abbreviations for Hawaiian structural genomes designated by Carr and Kyhos (1986) and Carr (2003), if documented, are indicated before taxon names. Subtribes of Madieae (Baldwin et al. 2002) and informal lineages of subtribe Madiinae (Baldwin 2003a, b) are indicated.

**Fig. 9.**
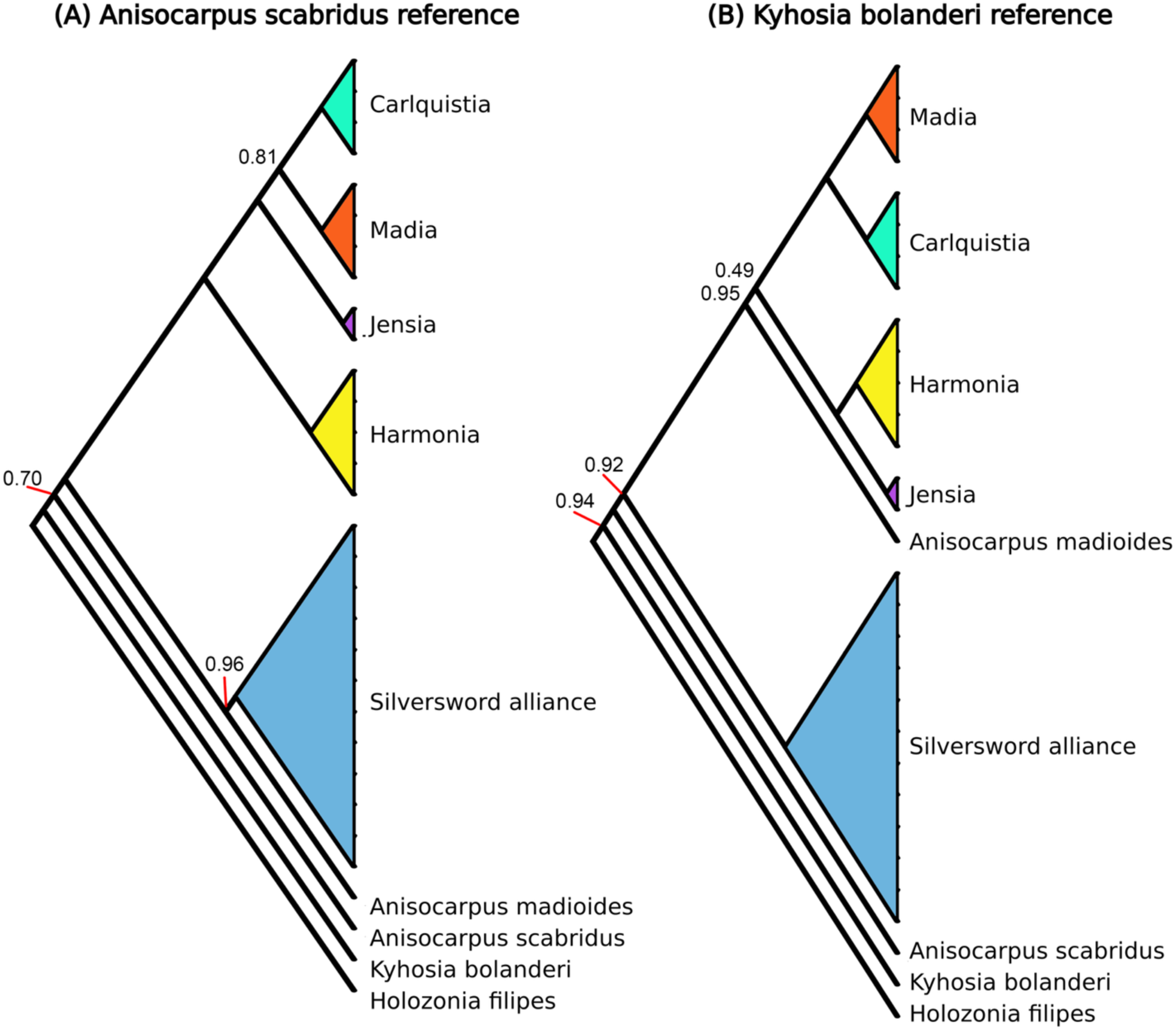
Collapsed summary trees showing two main patterns recovered for relationships of sequences of the Hawaiian silversword alliance when phased to tarweed references other than *Anisocarpus madioides.* Only taxa in the “Madia” lineage and its sister lineage, *Holozonia*, are shown, from multispecies coalescent (ASTRAL) trees of nuclear supercontig assemblies with silversword alliance reads phased to (A) *Anisocarpus scabridus*, when paired with *A. madioides*, and (B) *Kyhosia bolanderi*, when paired with *A. madioides.* Local posterior probabilities are shown at tree nodes if < 1.0.

### Phenotypic trait mapping

Trait mapping onto Madieae trees indicates an ancestrally perennial condition in the tribe, likely retained in Venegasiinae, Arnicinae, and Madiinae (in *Adenothamnus* and *Raillardella*) and more equivocally retained (possibly secondary) in *Constancea* B.G.Baldwin (Baeriinae) and *Hulsea* Torr. & A.Gray (Hulseinae), with widespread homology of the annual habit within Baeriinae and within much of Madiinae (Figure 10; Appendices S1A, S2A). Separate transitions from annual to perennial habit were resolved as likely in Baeriinae (e.g., in *Baeriopsis* and in *Eriophyllum*) and Madiinae (e.g., in *Centromadia perennis*). The “Madia” lineage was resolved as likely ancestrally (and secondarily) perennial, with at least one transition to the annual habit in the clade containing *Carlquistia, Harmonia*, *Jensia*, and *Madia*. Disk pappus was resolved as likely ancestral in tribe Madieae, with separate losses of pappus in Arnicinae (in *Arnica dealbata*), Baeriinae (in *Monolopia* DC. and *Pseudobahia* Rydb.), Madiinae (in *Centromadia pungens*, *Lagophylla* Nutt., *Hemizonia* DC., *Holocarpha* Greene, and *Madia*), and Venegasiinae (in *Venegasia* DC.) (Figure 11; Appendices S1B, S2B). Disk florets were resolved as likely ancestrally bisexual in tribe Madieae and in each of the five subtribes, with a loss of pistillate function in Arnicinae (in *Arnica dealbata*) and a complex history of transitions in Madiinae (Figure 12; Appendices S1C, S2C). Self-incompatibility was resolved as likely ancestral in tribe Madieae, with multiple losses, in Baeriinae (e.g., in *Lasthenia* Cass.) and Madiinae (e.g., in *Achyrachaena*, *Anisocarpus madioides*, *Jensia yosemitana*, *Lagophylla ramosissima*, *Madia*, and *Dubautia* sect. *Railliardia*), but evidently retained as the ancestral condition in the silversword alliance (Figure 13; Appendices S1D, S2D). Ray florets were resolved as likely ancestral in tribe Madieae, with multiple independent losses in Madiinae, i.e., in *Carlquistia*, *Raillardella* (A.Gray) Benth. & Hook.f., and the Hawaiian silversword alliance (Figure 14; Appendices S1E, S2E). Leaf dissection was resolved as evolving repeatedly in Baeriinae and Madiinae, with tarweed examples of evolution of deep leaf lobing in both the “Calycadenia” and “Layia” lineages but not in the clade encompassing the “Hemizonia” and “Madia” lineages and *Holozonia* (Figure 15; Appendices S1F, S2F).

**Fig. 10.**
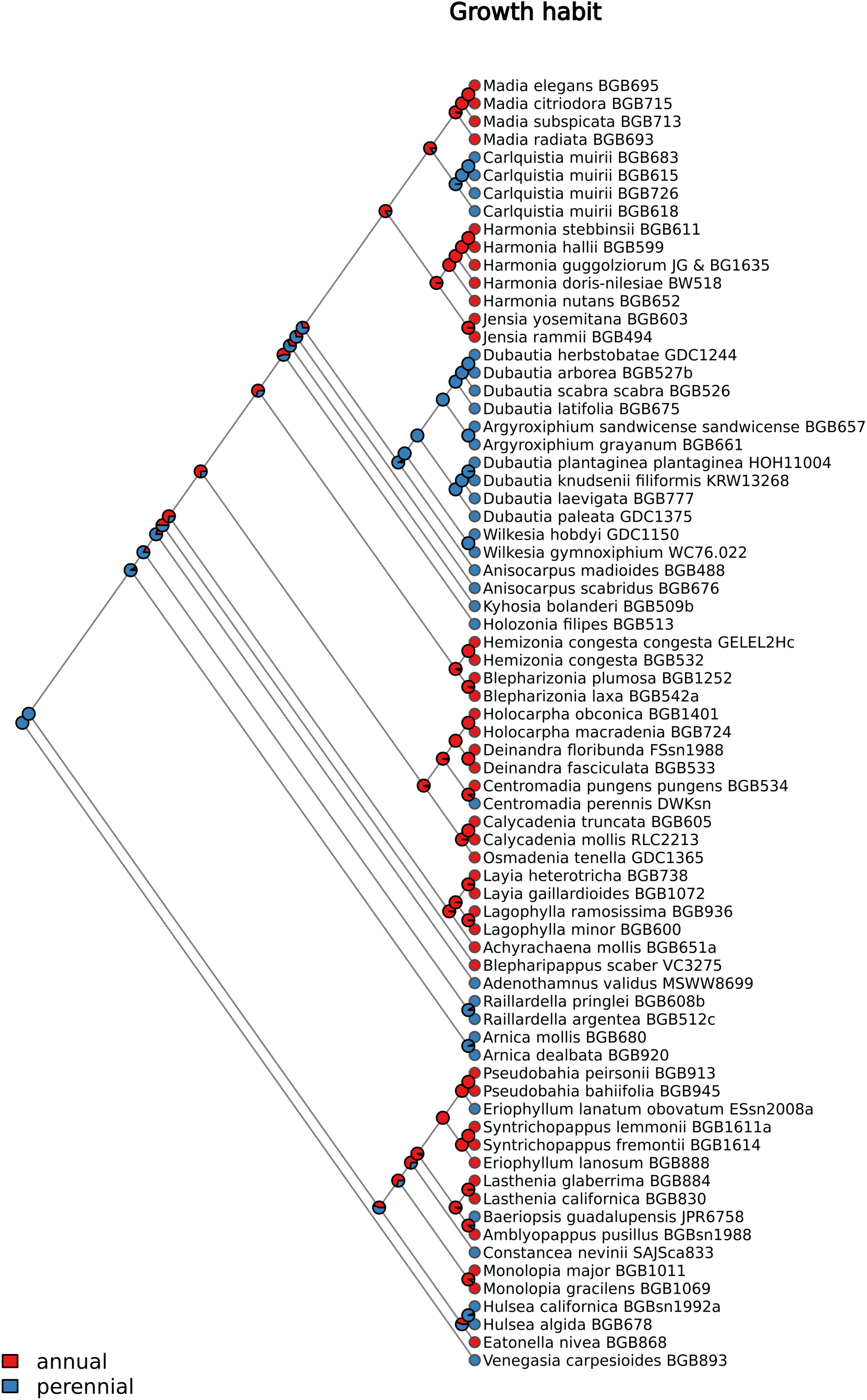
Maximum-likelihood mapping of growth habit onto an ultrametric tree with the multispecies coalescent (ASTRAL) topology in Figure 8.

**Fig. 11.**
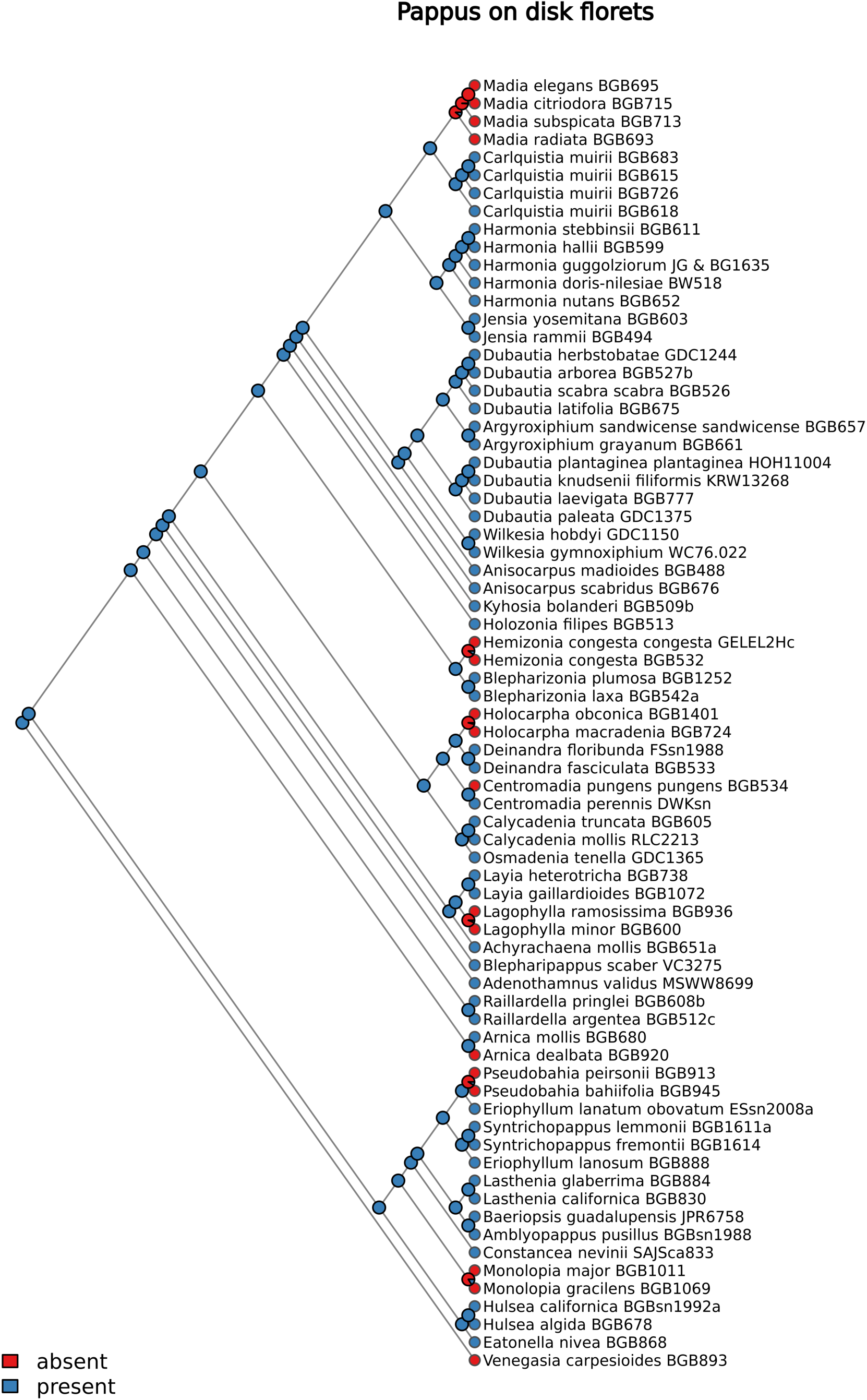
Maximum-likelihood mapping of pappus presence in disk florets onto an ultrametric tree with the multispecies coalescent (ASTRAL) topology in Figure 8.

**Fig. 12.**
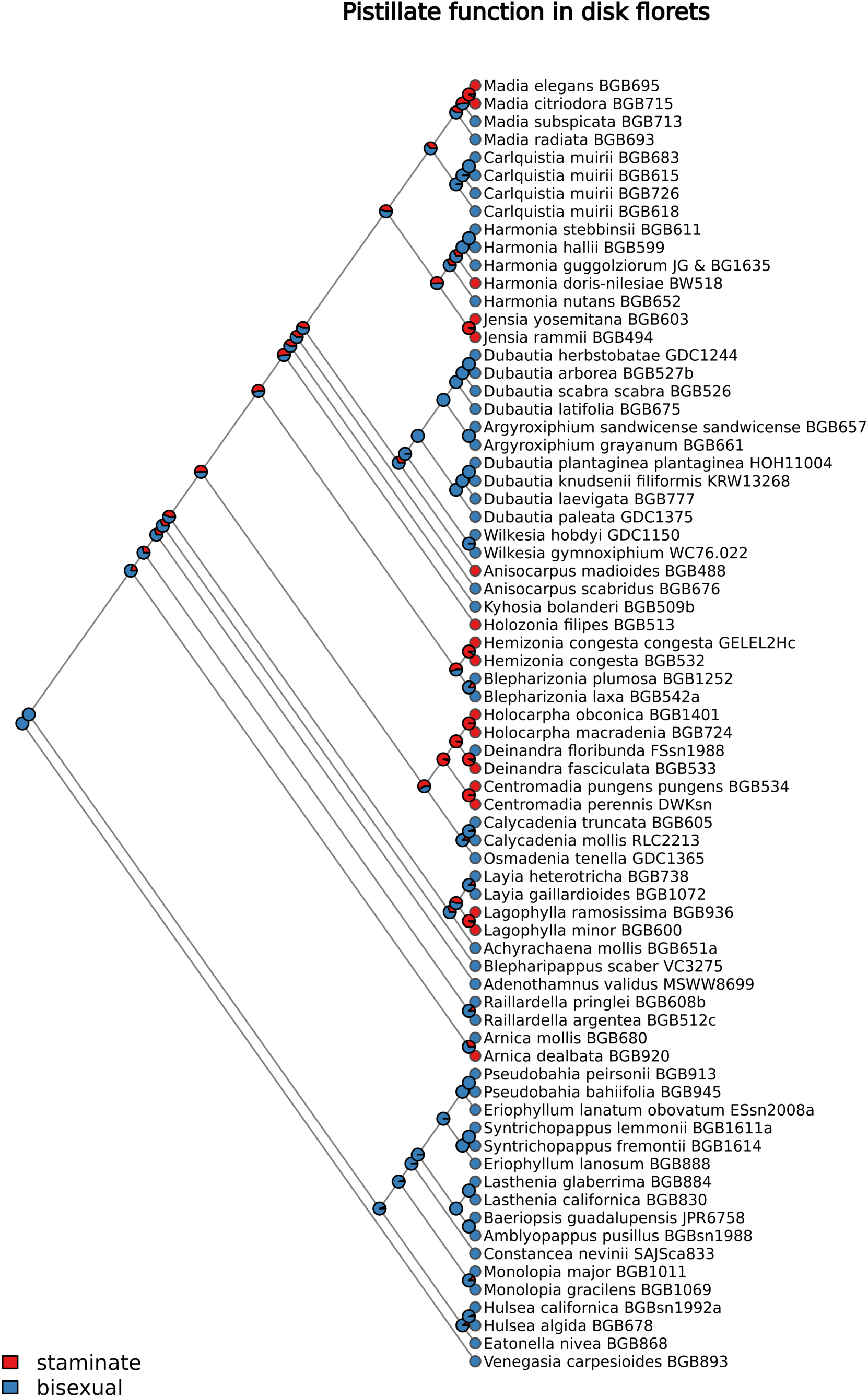
Maximum-likelihood mapping of pistillate function in disk florets onto an ultrametric tree with the multispecies coalescent (ASTRAL) topology in Figure 8.

**Fig. 13.**
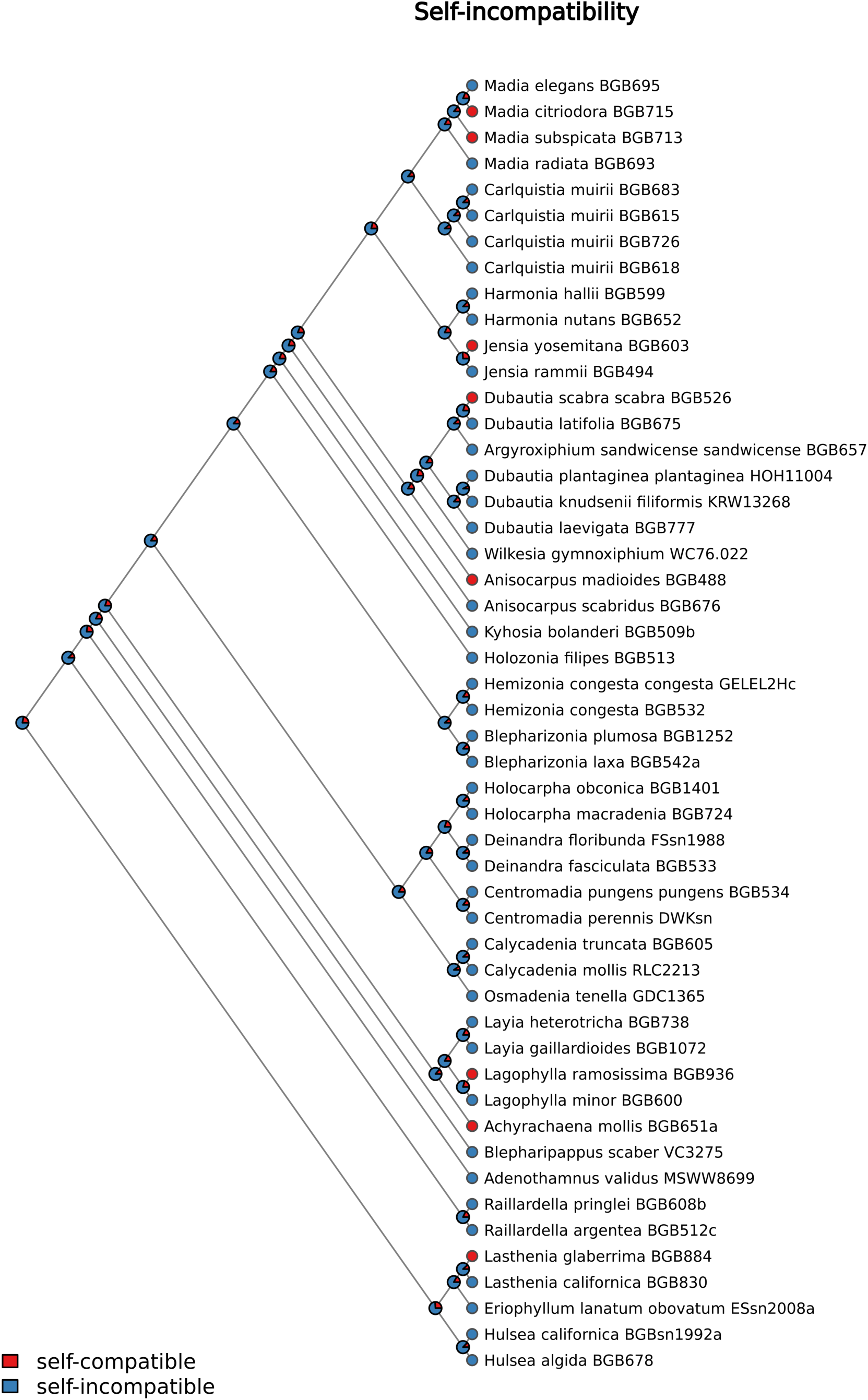
Maximum-likelihood mapping of self-incompatibility onto an ultrametric tree with the multispecies coalescent (ASTRAL) topology in Figure 8.

**Fig. 14.**
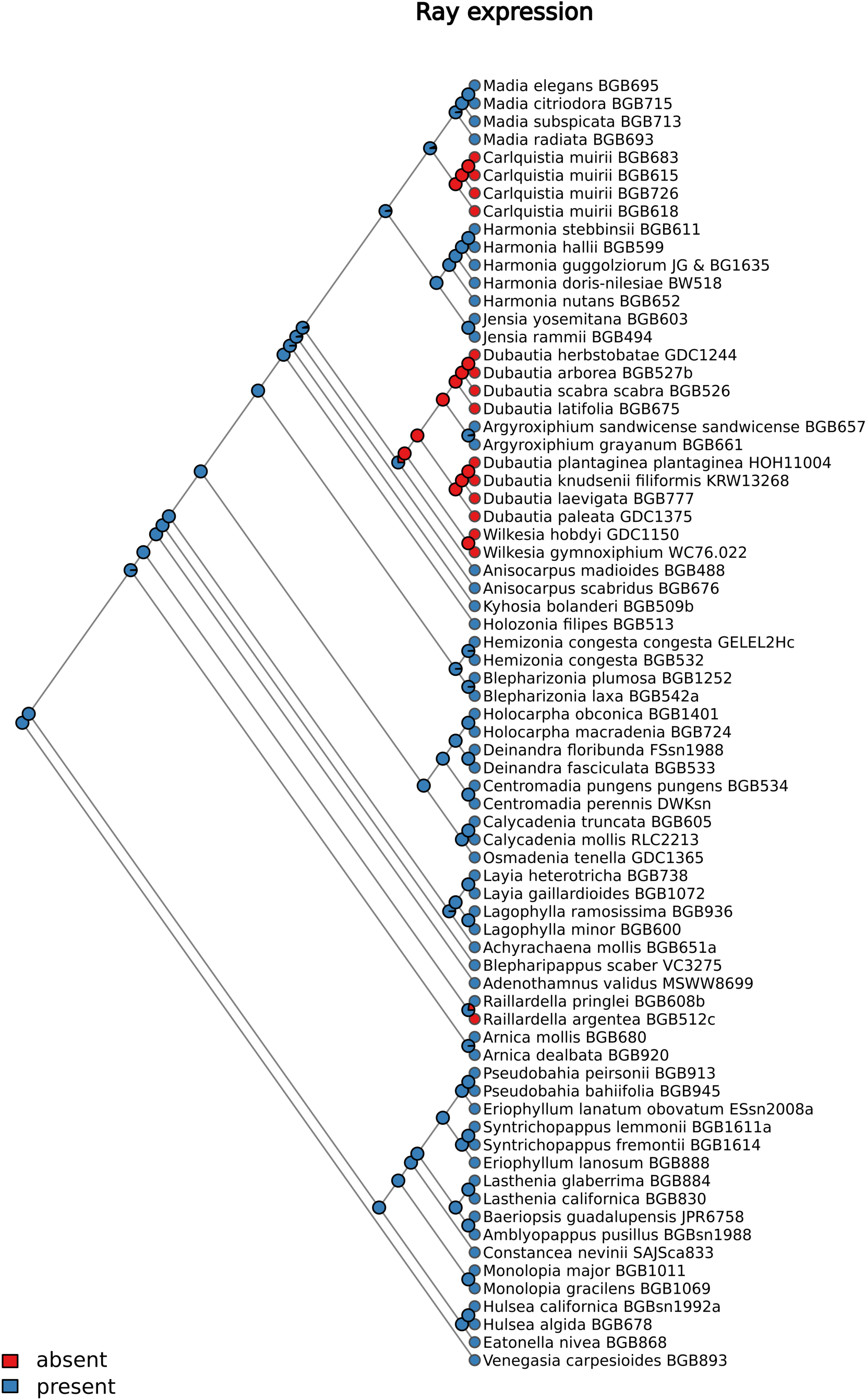
Maximum-likelihood mapping of ray floret presence onto an ultrametric tree with the multispecies coalescent (ASTRAL) topology in Figure 8.

**Fig. 15.**
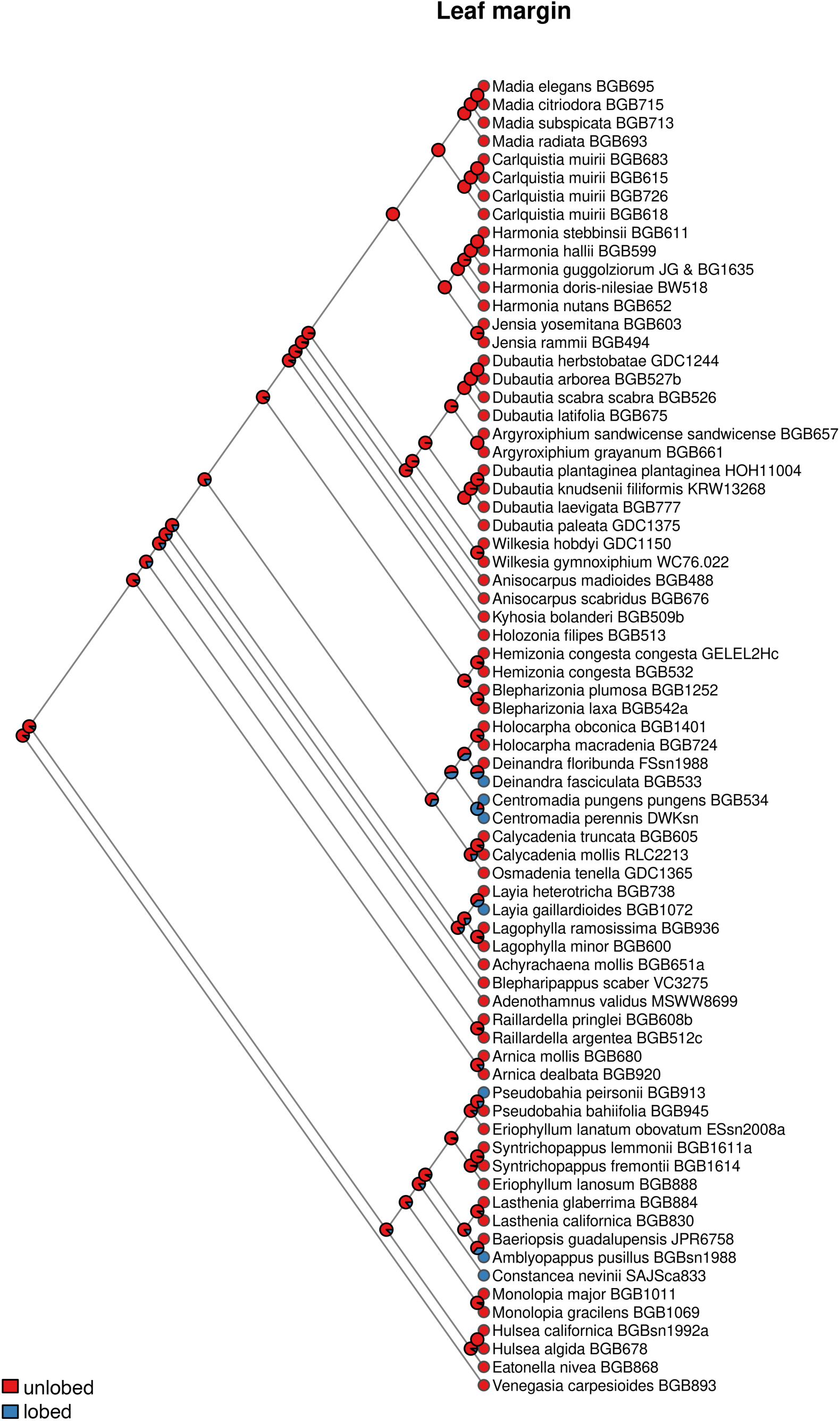
Maximum-likelihood mapping of leaf margin dissection onto an ultrametric tree with the multispecies coalescent (ASTRAL) topology in Figure 8.

### Chromosome-number evolution

Likelihood-based inference of chromosome-number evolution in the “Madia” lineage (Appendix S3A--C) indicated a highly probable (0.99) ancestral condition of 2*n* = 7_II_ for the most recent common ancestor of the silversword alliance and its North American sister group based on all of the trees analyzed. The same ancestral condition of 2*n* = 7_II_ was indicated with similarly high probability (0.97--0.99) for all deeper nodes in the “Madia” lineage, including its root node. An ancestral chromosome number of 2*n* = 14_II_ was inferred as highly probable (0.98--0.99) for the silversword alliance, with descending dysploidy within *Dubautia* sect. *Railliardia* to 2*n* = 13_II_, which was the best supported (0.91--0.94) condition for the most recent common ancestor of *D. arborea* and *D. herbstobatae*.

## DISCUSSION

### Allopolyploidy of the silversword alliance

Successful phasing of sequencing reads to different subgenomes of the silversword alliance (Figures 1, 2, and 7A--C) using diploid North American tarweed representatives of distinct sublineages of the “*Madia*” lineage (Figures 3 and 4A, D, E) reinforces earlier evidence of allopolyploidy of the Hawaiian radiation (Barrier et al., 1999), now with genome-scale data. Ancestral allopolyploidy of two other major insular radiations, the Galápagos giant daisies (*Scalesia* Arn.) and the Hawaiian mints (*Haplostachys* Hillebr., *Phyllostegia* Benth., *Stenogyne* Benth.) also have been recently confirmed with genomic evidence (Cerca et al., 2022; Tomlin et al., 2024). Greater genomic variation and fixed heterozygosity in allopolyploids, as opposed to autopolyploids, may promote colonization and evolutionary success on remote oceanic islands by limiting loss of genetic variation associated with founder effects and by providing an intrinsic source of variation for adaptation (see Soltis and Soltis, 2000), e.g., by changes in gene expression or genome organization (see Barker et al., 2016a). The relative frequency of ancestral allo- vs. autopolyploidy in island plants, however, has been unclear (Meudt et al., 2021).

Evidence from both clade association of tetraploid Hawaiian taxa reads (Figures 5 and 6) and subsequent phylogenomic analyses (Figures 7A--C and 8) that one subgenome of the silversword alliance (hereafter, Subgenome A) is most closely related to a single extant North American species, *Anisocarpus madioides* (Figure 3), indicates an even more intimate relationship of the Hawaiian radiation to modern tarweeds than previously resolved. Results of earlier phylogenetic analyses of low-copy nuclear genes led to the conclusion that *Anisocarpus* may be most closely related to one of the two Hawaiian subgenomes (Barrier et al., 1999, 2001; Remington and Purugganan, 2002), but those studies included *A. madioides* or *A. scabridus* (Figure 4A) — not both — based on nrDNA evidence that the two species of *Anisocarpus* constituted a clade (Baldwin, 1996; Baldwin and Sanderson, 1998). Instead, the nuclear phylogenomic data strongly resolve *Anisocarpus* as a paraphyletic grade within the “Madia” lineage and the only example of non-monophyly among diploids of currently recognized continental genera in subtribe Madiinae (Figure 8).

The precise relationship of the other Hawaiian subgenome (hereafter, Subgenome B) to modern diploid tarweeds (Figure 9) is less conclusive than the relationship of Subgenome A to *Anisocarpus madioides.* These results suggest that the mainland ancestor of Subgenome B belonged to a lineage that diverged from the backbone of the “Madia” lineage prior to divergence of the *A. madioides* lineage from a common ancestor with the lineage that gave rise to *Carlquistia, Harmonia, Jensia,* and *Madia*, or that split off after that divergence event, e.g., in the early history of the *A. madioides* lineage. Previous phylogenetic analyses of low-copy nuclear genes (Barrier et al., 1999, 2001; Remington and Purugganan, 2002; Purugganan et al., 2003) may approximate these findings, although differences in taxon sampling complicate comparisons. The possibility that uncertainty in the relationship of Subgenome B may in part reflect post-polyploidization processes, such as homeologous exchange and subgenome dominance, remains to be explored.

### Ancestral characteristics of the silversword alliance

This new perspective on relationships of the silversword alliance allows for insights into the characteristics of the colonizing ancestor based on conditions shared by the closest living relatives of subgenome lineages. *Anisocarpus*, as delimited to include *A. madioides* (Figure 3) and *A. scabridus* (Figure 4A), now appears to be a genus united by states that diagnose a grade, rather than a clade, along which the lineages corresponding to the two subgenomes of the silversword alliance appear to have diverged. Those states include the perennial habit, extensively leafy stems, radiate heads, ellipsoid to spheric involucres, yellow anthers, well-developed disk pappus, and 2*n* = 7_II_ (Baldwin, 2003b). *Anisocarpus madioides* and *A. scabridus* are the only members of the “Madia” lineage and the only perennials in the tarweed subtribe, Madiinae, with seven pairs of chromosomes, in keeping with the ancestral 14-paired chromosomal condition in the silversword alliance (Carr and Kyhos, 1981; Witter and Carr, 1988; Baldwin et al., 1990, 2021; Carr, 2003). Radiate heads are universal among continental members of the “Madia” lineage, but are present only in the true silverswords and greenswords (*Argyroxiphium*; Figure 1A, B) among the Hawaiian taxa, with *Dubautia* (Figures 1E--I and 2) and *Wilkesia* (Figure 1C, D) being strictly discoid-headed and even some members of *Argyroxiphium* having limited ray expression, e.g., *A. grayanum* (Figure 1A; see Carr, 1985). *Anisocarpus scabridus* (Figure 4A) also has irregular expression of ray florets, with only 1--3 rays per head and some heads discoid, which is unusual variation within a continental tarweed (Baldwin, 2003b).

Ecologically, the combination of traits that were inferred by ML to be ancestral in the silversword alliance includes some favorable for long-distance dispersal and colonization. The perennial habit is uncommon among continental tarweeds but was estimated by ML to be ancestral in the “Madia” lineage and in the silversword alliance (Figure 10; Appendices S1A, S2A), as was self-incompatibility (Figure 13; Appendices S1D, S2D). The ability of a perennial colonist to persist vegetatively after long-distance dispersal has been suggested as potentially critical to establishment of a putatively self-incompatible ancestor of the silversword alliance facing an S-allele bottleneck following dispersal to the Hawaiian Islands (Carr et al., 1986). The presence of pappus elements on the disk fruits, which can aid in attachment of fruits to feathers or fur (Carlquist, 1974), also was estimated as ancestral in the silversword alliance, with subsequent loss of pappus in the continental genus *Madia* (Figure 11; Appendices S1B, S2B). Bisexual expression in disk florets, which allows formation of disk fruit, as opposed to disk florets being merely staminate in function, was also estimated as ancestral in the silversword alliance (Figure 12; Appendices S1C, S2C). Ray florets, which in subtribe Madiinae produce fruit enveloped by sticky involucral bracts that readily attach to animals (Carlquist, 1966), were estimated as ancestral in tribe Madieae, with isolated losses of ray florets and their associated involucral bracts (e.g., in *Carlquistia*, *Raillardella*, and the silversword alliance). The estimated absence of ray florets in the most recent common ancestor of the silversword alliance (Figure 14; Appendices S1E, S2E) warrants cautious interpretation, given remaining uncertainty in resolution of relationships among the four main Hawaiian sublineages and potentially heightened selection for loss of ray florets and associated dispersibility in the insular environment (Carlquist, 1974).

The morphology, ecology, and biogeography of *Anisocarpus madioides* (Figure 3), the species strongly resolved as most closely related to Subgenome A of the silversword alliance (Figures 7A--C and 8), align well with expectations for high dispersibility, including some traits estimated by ML to be ancestral in the Hawaiian lineage. *Anisocarpus madioides* is a rhizomatous perennial and by far the most geographically widespread perennial in the tarweed subtribe Madiinae, with a distribution extending north from the Peninsular Ranges of southwestern California to southern British Columbia, from coastal to montane woodland habitats (Baldwin, 2003b). Ray fruits of *A. madioides* are tightly enveloped by sticky involucral bracts that are readily adhesive (Figure 3D) and may help explain major disjunctions in the distributional range of the species, especially among southern occurrences. Colonization ability of *A. madioides* is also evident from its higher density along trails and in disturbances or somewhat open sites in woodlands and forests.

The chromosomal condition in *Anisocarpus madioides* of 2*n* = 7_II_, estimated with high probability to be shared by the most recent common ancestor of tarweeds and silverswords and with more distant ancestors in the “Madia” lineage (Appendix S3A--C), not only strengthens the hypothesis of homology between the chromosome numbers of *A. madioides* and Subgenome A of the silversword alliance but also indicates that the diploid ancestor of Subgenome B probably shared that chromosome number, as well. Chromosome-number evolution was studied from trees with silversword alliance sequences assembled from reads phased to different, opposing references, in an effort to examine histories of both subgenomes.

Characteristics of *A. madioides* that were not estimated to be ancestral in the silversword alliance include self-compatibility (Figure 13; Appendices S1D, S2D) and functionally staminate disk florets (Figure 12; Appendices S1C, S2C). Together, these two traits operate to promote outcrossing but permit selfing, with all florets unisexual (and a brief period when ray floret stigmatic areas are receptive prior to adjacent disk florets opening and presenting pollen) but with capacity for self-fertilization by pollen transfer from disk to ray florets if cross-pollination fails to occur. Showiness of heads of *A. madioides* runs counter to the expectation of reduced rays for the selfing syndrome in Compositae (Ornduff, 1969; Sicard and Lenhard, 2011) but is consistent with results from ML trait mapping that the shift from self-incompatibility to self-compatibility occurred relatively recently, after divergence of the *A. madioides* lineage (Figure 13; Appendices S1D, S2D). Involvement of a diploid parent with self-compatible and functionally staminate disk florets in formation of the allopolyploid lineage that gave rise to the silversword alliance is not necessarily in conflict with widespread self-incompatibility and ubiquitously bisexual disk florets throughout the Hawaiian radiation. Synthetic allopolyploids produced from hybrids between self-compatible and self-incompatible species of *Senecio*, although strictly self-compatible in the F_1_ generation, included ca. 7--9% self-incompatible individuals in the F_2_ generation (Brennan and Hiscock, 2010), a result consistent with dominance relationships among S-alleles in the sporophytic self-incompatibility system of Compositae (see Hiscock, 2000; Crawford et al., 2024). Also, Clausen et al. (1945), in generating synthetic allopolyploids between members of the “Madia” lineage, found that bisexual function of disk florets was dominant to the functionally staminate condition in both examples of such hybrid polyploid lines, in *Harmonia nutans* (bisexual) × *Jensia rammii* (staminate) and in *Madia gracilis* (bisexual) × *M. citriodora* (staminate).

### Biological prospects for allopolyploidization

The biological potential for allopolyploidization involving perennial diploid members of the “Madia” lineage has been demonstrated previously, with production of vigorous but nearly sterile F_1_ hybrids between *Carlquistia muirii* and *Kyhosia bolanderi* (Kyhos et al., 1990; Carr et al., 1996) and between *Anisocarpus scabridus* and *Carlquistia muirii* (Barrier et al., 1999). These synthetic hybrids showed limited to no chromosomal association at meiosis and produced a low percentage of stainable pollen, dominated by large, tetraporate (rather than normal, triporate) grains that represent an unreduced, diploid condition (Carr et al., 1996). Artificial hybrids between the two taxa in the “Madia” lineage with 2*n* = 7_II_, *Anisocarpus madioides* [as pistillate parent; *Baldwin 845* (JEPS)] and *A. scabridus* [as staminate parent; *Baldwin 620* (JEPS)], have been subsequently produced, with similar meiotic chromosomal behavior and similarly low pollen stainability (5--12.4%; *x̅* = 8.7%), again with stainability essentially confined to large, tetraporate grains.

Although synthetic allopolyploids were not produced from any of these F_1_ hybrids, Carr et al. (1996) showed that the stainable tetraporate pollen produced by such hybrids (e.g., by *Kyhosia bolanderi* × *Carlquistia muirii*) was unreduced and effective to produce tetraploid hybrids with a species of the Hawaiian silversword alliance, *Dubautia scabra*, that served as pistillate parent. Highly irregular and reduced meiotic chromosomal pairing in these synthetic tetraploid tarweed--silversword hybrids and in additional, triploid tarweed--silversword hybrids, between diploid perennials in the “Madia” lineage and members of the silversword alliance, did not allow for conclusions about chromosomal homology between mainland and Hawaiian Madiinae (Carr et al., 1996). The diploid hybrids do indicate that despite ca. 5 million years of evolutionary divergence since formation of the allopolyploid silversword alliance lineage (Landis et al., 2018), the potential remains for producing hybrids with viable unreduced pollen (and presumably some percentage of unreduced ovules) between continental perennials in the “Madia” lineage that could be effective for producing allopolyploid progeny.

### Relationships within the silversword alliance

Based on the unrooted multispecies coalescent trees from the three phasings involving *Anisocarpus madioides* as one of the two diploid tarweeds used as references (Figures 7A--C), four major clades are well-supported within the Hawaiian radiation that correspond to the two rosette-plant genera, *Argyroxiphium* (Figure 1A, B) and *Wilkesia* (Figure 1C, D), and two major clades within *Dubautia* in the broad sense, i.e., (1) *D*. sect. *Dubautia* sensu G.D. Carr (1985) (Figure 2) and (2) *D. latifolia* + *D.* sect. *Railliardia* sensu G.D. Carr (1985) (Figure 1E--I). If the tree from the fourth phasing is rooted along the branch corresponding to the internal branch connecting the two subgenome clades in Figures 7A--C, then it, too, resolves the same four major groups (Figure 7D).

The two major *Dubautia* clades, which are not resolved here as sister groups, are morphologically and chromosomally distinct (Carr, 1985, 2003; Carr and Kyhos, 1986) and correspond to groups treated by Gray (1861), and later Hillebrand (1888), Sherff (1935), Skottsberg (1944), Degener and Degener (1958), and St. John (1981), as distinct genera, *Dubautia* and *Railliardia* Gaudich. Morphologically, *Railliardia* sensu A. Gray (Figure 1E--I) differs most conspicuously from *Dubautia* sensu A. Gray (Figure 2) by its plumose rather than merely ciliate, fimbriate, or barbellate disk pappus elements, and more generally by its consistently cup-like false-involucre of basally fused bracts, rather than a more variably shaped false-involucre of often free bracts, sometimes with internal paleae, as noted by Gray (1861). Chromosomally, members of *Dubautia* sensu A. Gray have 2*n* = 14_II_ (as do *Argyroxiphium* and *Wilkesia*) and those that have been characterized cytogenetically have one of three structural arrangements of the nuclear genome known only from these core members of *Dubautia*: Dubautia Genome 1, 2, or 3 (Carr and Kyhos, 1986; Carr, 2003). Members of *Railliardia* sensu A. Gray have 2*n* = 14_II_ or 13_II_, and are characterized by having one of two nuclear structural arrangements known only from *Railliardia* sensu A. Gray: Dubautia Genome 4 or 5 (Carr and Kyhos, 1986; Carr, 2003). Representatives of the diverse clade with 13, rather than 14, pairs of chromosomes (represented here by *D. arborea* and *D. herbstobatae*) share a common genomic structural arrangement (Dubautia Genome 5) and are consistently nested here within a grade of two species (*D. latifolia* and *D. scabra*) with a 14-paired chromosomal arrangement (Dubautia Genome 4) that can be inferred to have given rise to the 13- paired condition by a Robertsonian translocation, i.e., one reciprocal translocation between two chromosomes with small heterochromatic arms and loss of the resulting centric fragment (Carr and Kyhos, 1986).

Biogeographically and ecologically, *Dubautia* sensu A. Gray (Figure 2) and *Railliardia* sensu A. Gray (Figure 1E--I) are also somewhat distinct. All species of *Dubautia* sensu A. Gray are native to the oldest high island, Kauài, and all but two are endemic there, whereas *Railliardia* sensu A. Gray is primarily found on the other, younger high islands of the Hawaiian chain, with only *D. latifolia* on Kauài. *Dubautia* sensu A. Gray comprises taxa known from wet or mesic habitats, whereas *Railliardia* sensu A. Gray also comprises taxa that are restricted to dry habitats (Carr, 1985).

Consistent resolution here of *Dubautia latifolia* (Figure 1E) as sister to other taxa of *Railliardia* sensu A. Gray (Figures 1F--I, 7 and 8) helps to resolve long-standing uncertainty about the relationships of arguably the most distinctive species in the silversword alliance. *Dubautia latifolia* is unique within the Hawaiian radiation (and tribe Madieae) as a woody vine or liana with broad, petiolate leaves of distinctive isodiametric reticulate venation, rather than having parallel or parallel-trending venation patterns, as in other members of the silversword alliance. Gray (1861), Sherff (1935), and Carr (1985) treated *D. latifolia* as warranting recognition as a separate taxonomic section. Enzyme electrophoretic data showed *D. latifolia* to be highly divergent from other members of the silversword alliance (Witter and Carr, 1988). Subsequent phylogenetic studies indicated incongruent relationships of *D. latifolia.* Chloroplast DNA and nuclear ribosomal DNA trees, for example, placed *D. latifolia* well apart from *D. scabra* (Figure 1F), with which it shares an otherwise unique genomic arrangement (Carr, 2003), as noted above, and from taxa of *D.* sect. *Railliardia* (Figure 1F--I) in general (Baldwin et al., 1990, 2021; Baldwin and Robichaux, 1995). Those data also strongly supported a nested position of *D. scabra* among the taxa with 13 pairs of chromosomes, in conflict with cytogenetic evidence of relationships (Carr and Kyhos, 1986). Phylogenetic analysis of floral homeotic genes, by contrast, resolved a moderately supported clade uniting representatives of *Railliardia* sensu A. Gray, with *D. latifolia* and *D. scabra* constituting a clade sister to the sole representative of the taxa with 13 pairs of chromosomes, *D. sherffiana* (Barrier et al., 1999). A later study of floral homeotic genes, including more representatives of the 13-paired taxa, placed *D. scabra* just outside a 13-paired clade, but did not sample *D. latifolia* (Friar et al., 2008). The phylogenomic data come closest to approximating the floral homeotic gene results and, as noted above, are fully congruent with cytogenetic evidence that *D. latifolia* is a biogeographic link back to the oldest high island, Kauài, for the younger-island endemic radiation of *Dubautia* sect. *Railliardia*.

### Evolutionary and ecological patterns across tribe Madieae

Although finer- scale analyses of phylogenomic relationships among members of Madieae will be published elsewhere by the authors, the broad taxon sampling here yielded some insights into general evolutionary patterns. Trait mapping corroborates a perennial ancestry of this annual-rich, primarily Californian tribe (Figure 10; Appendices S1A, S2A), with ML support for homology of the perennial habit among most or all of those perennials with 2*n* = 19_II_. Those findings align with interpretation of the widespread annual habit and generally lower chromosome numbers in Madieae as evolutionarily derived, associated with xeric conditions of low-elevation, generally interior habitats (Baldwin and Wessa, 2000). Stebbins (1950) predicted such descending dysploidy in annuals with selection for reduced recombination under an exacting, dry environment. Robust placements of *Arnica* as sister to tarweeds and, in turn, of *Raillardella* as sister to other tarweeds (Figure 8) reinforce the interpretation that the rhizomatous perennial condition, high base chromosome number, montane ecology, and other *Arnica*-like traits of *Raillardella* are probably ancestral in Madiinae despite the rarity of those conditions in the tarweed subtribe (Baldwin and Wessa, 2000; Baldwin, 2003a). Secondary reacquisition of the perennial habit in Madieae was generally mapped to lineages associated with maritime (including insular), otherwise mesic, or montane settings, where selection for persistence may be expected (Carlquist, 1974; Drummond, 2008; see Baldwin, 2014).

The tribal-scale mapping of leaf margin evolution in Madieae (Figure 15; Appendices S1F, S2F) provides a novel perspective on the extraordinary diversity in leaf morphology and anatomy that arose during adaptive radiation of the silversword alliance. Leaf shape and venation patterns across the silversword alliance span an exceptional range, including sessile, strap-shaped leaves with parallel venation and few cross-connecting veinlets (in *Wilkesia*; Figure 1C, D), like those of a graminoid monocot, to petiolate, oblong-elliptic leaves with highly reticulate venation (in *Dubautia latifolia*; Figure 1E), like those of a typical mesophytic eudicot (Carr, 1985; Blonder et al., 2016). Lack of leaf dissection in the silversword alliance, however, is a sharp contrast with leaf evolution in *Scalesia*, another example of insular adaptive radiation in the Heliantheae alliance and the most diverse plant lineage in the Galápagos. In *Scalesia*, repeated evolution of leaf dissection has been resolved, putatively associated with evolution to warm, dry habitats, where dissected leaves may aid in heat dissipation and transpiration reduction (Fernández-Mazuecos et al., 2020; Bieker et al., 2026).

Trait mapping indicates that lack of leaf dissection in the silversword alliance extends to the rest of the “Madia” lineage and to the deeper lineage also including *Holozonia* and the “Hemizonia” lineage, whereas deeply lobed leaves have evolved repeatedly elsewhere in tribe Madieae, including in both the “Calycadenia” and “Layia” lineages of the tarweed subtribe Madiinae (Figures 8 and 15; Appendices S1F, S2F). In *Layia gaillardioides*, for example, Clausen (1951) documented distinct ecotypes that varied in degree of heritable leaf lobing, with greater lobing in those associated with hotter, drier environments (see Baldwin, 2006). Whether absence of leaf dissection in the deep lineage that includes the silversword alliance and closely related tarweeds reflects anatomical or physiological innovation or constraint remains to be determined, but has not prevented these extraordinary plants from diversifying into dry habitats in both California and the Hawaiian Islands.

### Taxonomic implications

Phylogenomic results corroborate the subtribal taxonomy of Madieae (Figure 8) proposed earlier (Baldwin et al., 2002) and relationships among subtribes resolved by Zhang et al. (2021) based on more limited taxon sampling. Relationships among genera resolved here also support the four main clades in the tarweed subtribe Madiinae discussed previously (Baldwin, 2003a, b), i.e., the “Calycadenia”, “Hemizonia”, “Layia”, and “Madia” lineages, the last including both California tarweeds and the Hawaiian silversword alliance (Figure 8). Within the “Madia” lineage, one of the main findings here, that *Anisocarpus madioides* (Figure 3) is more closely related to one of the two subgenomes of the silversword alliance than it is to its congener, *A. scabridus* (Figure 4A), warrants return of *A. scabridus* to *Raillardiopsis* Rydb., as *R. scabrida*. Evident taxonomic problems in *Dubautia* and *Eriophyllum* Lag. (Figure 8) will be addressed by the authors elsewhere, with presentation of more detailed phylogenomic data across continental Madieae and the Hawaiian silversword alliance.

## CONCLUSIONS

The ability to phase sequencing reads to subgenome using diploid tarweed references (Figures 7A--C) allowed for nuclear phylogenomic analysis of the allotetraploid silversword alliance without a reference genome. Fortunately, an extant North American tarweed, *Anisocarpus madioides* (Figure 3), is closely related to one of the two subgenomes (Subgenome A) of the silversword alliance, and other North American members of the “Madia” lineage (Figure 4A, D, E) retain enough similarity to the other silversword alliance subgenome to be effective Subgenome B references, when paired with *A. madioides*. Advances in understanding subgenomic relationships in the silversword alliance have importance for more than phylogenomics. As reference genomes are developed for the silversword alliance, understanding the closest living relatives of each subgenome should facilitate resolution of post-polyploidization molecular and chromosomal evolution and allow more nuanced perspectives on the genomic underpinnings of one of earth’s most spectacular adaptive radiations.

## ACKNOWLEDGMENTS

The authors thank the National Tropical Botanical Garden, National Science Foundation (DEB-9458237, DEB-2040347, DEB-2040081, DEB-2040097), Lawrence R. Heckard Endowment Fund of the Jepson Herbarium, and the late Roderic B. Park and other generous Friends of the Jepson Herbarium for research support; Michael S. Barker and Robert H. Robichaux for collaboration on the unpublished silversword alliance transcriptomes used for bait design in this study; Lowell Ahart, Susan J. Bainbridge, Gerald D. Carr, Robert L. Carr, the late Winona P. Char, Victor P. Claassen, Gregory English-Loeb, the late Jack and Betty Guggolz, the late Robert W. Hobdy, Nick Jensen, Steven A. Junak, the late Donald W. Kyhos, Henry Oppenheimer, the late Vernon H. Oswald, Steven Perlman, Jon P. Rebman, Robert H. Robichaux, Erin Shapiro, Fred T. Sproul, the late Lani Stemmermann, the late Christopher H. Thayer, Barbara Williams, Martha S. Witter, and Kenneth R. Wood for collection assistance; the National Park Service, U.S. Forest Service, Hawaii Department of Forestry and Wildlife, and Hawaii State Parks for research or collection permits; UC Berkeley Research IT for use of the Savio High Performance Computing Cluster; Matthew Berger, Gerald D. Carr, Preston Ernest, Dean Lyons, C. Newfield, Jen Pagel, Zach Pezzillo, Kenneth R. Wood, and Jacob (earthbound_adventurer) for making their photos available by permission or through a Creative Commons License, and Madieae researchers, past and present, for their contributions, encouragement, and inspiration.

## AUTHOR CONTRIBUTIONS

All authors contributed to conception and design of the study, acquisition of data, and/or analysis and interpretation; BGB drafted the manuscript; SF prepared the figures; AK and SF executed most data analyses and management; BLW conducted lab work; WAF designed baits for target enrichment; and all authors approved the final version of the manuscript.

## DATA AVAILABILITY STATEMENT

Additional methodological details, scripts, and trees can be found at github.com/susanfawcett/Madieae/tree/Allopolyploid_Origin_Silverswords. Sequence alignments can be found in the DRYAD data repository (https://doi.org/10.5061/dryad.cjsxksnnr) and raw sequence data can be found in the NCBI sequence read archive (Bioproject PRJNA1497529). Additional supporting information may be found online in the Supporting Information section at the end of the article:

Appendix S1. Maximum-likelihood mapping of phenotypic traits in tribe Madieae based on an ultrametric tree with multispecies coalescent (ASTRAL) topology for nuclear supercontigs and silversword alliance reads phased to *Kyhosia bolanderi*, when paired with *Anisocarpus madioides*.

Appendix S2. Maximum-likelihood mapping of phenotypic traits in tribe Madieae based on an ultrametric tree with multispecies coalescent (ASTRAL) topology for nuclear supercontigs and silversword alliance reads phased to *Carlquistia muirii*, when paired with *Anisocarpus madioides*.

Appendix S3. Maximum-likelihood inference of ancestral chromosome numbers and polyploidization in the “Madia” lineage of the tarweed subtribe Madiinae, based on ultrametric trees with multispecies coalescent (ASTRAL) topologies for nuclear supercontigs and silversword alliance reads phased to (A) *Anisocarpus madioides*, when paired with *A. scabridus*, (B) *Kyhosia bolanderi*, when paired with *A. madioides*, and (C) *Carlquistia muirii*, when paired with *A. madioides*.

## APPENDICES

Appendix 1. Voucher information for samples included in this study. Abbreviations: *BGB* = *Bruce G. Baldwin* (and colleagues).

***Achyrachaena mollis*** Schauer: USA, California, Solano Co., *BGB 651* (DAV). ***Adenothamnus validus*** (Brandegee) D.D.Keck: MEXICO, Baja California, Municipio Ensenada, *Martha S. Witter W86-99* (DAV). ***Amblyopappus pusillus*** Hook. & Arn.: MEXICO, Baja California, Municipio Ensenada (Isla San Martín), *BGB s.n.*, 1988 (UC). ***Anisocarpus madioides*** Nutt.: USA, California, Napa Co., *BGB 488* (DAV). ***Anisocarpus scabridus*** (Eastw.) B.G.Baldwin [= ***Raillardiopsis scabrida*** (Eastw.) Rydb.]: USA, California, Lake Co., *BGB 676* (DAV). ***Argyroxiphium grayanum*** (Hillebr.) O.Deg.: USA, Hawaii, Maui (West Maui), *BGB 661* (DAV). ***Argyroxiphium sandwicense*** subsp. ***sandwicense*** DC.: USA, Hawaii, Hawaii Island, *BGB 657* (DAV). ***Arnica dealbata*** A.Gray: USA, California, Tehama Co., *BGB 920* (JEPS). ***Arnica mollis*** Torr. & A.Gray: USA, California, Alpine Co., *BGB 680* (DAV). ***Baeriopsis guadalupensis*** J.T.Howell: MEXICO, Baja California, Municipio Ensenada (Isla Guadalupe), *Jon P. Rebman 6758* (UC). ***Blepharipappus scaber*** Hook.: USA, California, Lassen Co., *Bob Powell & Vic Claassen 3275* (DAV). ***Blepharizonia laxa*** (A.Gray) Greene: USA, California, San Joaquin Co., *BGB 542a* (DAV). ***Blepharizonia plumosa*** (Kellogg) Greene: USA, California, Contra Costa Co., *BGB 1252* (JEPS). ***Calycadenia mollis*** A.Gray: USA, California, Madera Co., *Robert L. Carr 2213* (EWU). ***Calycadenia truncata*** DC.: USA, California, Tehama Co., *BGB 605* (DAV). ***Carlquistia muirii*** (A.Gray) B.G.Baldwin: USA, California, Fresno Co., *BGB 615* (DAV); Monterey Co., *BGB 618* (DAV); Tulare Co., *BGB 683* (DAV); *BGB 726* (DAV). ***Centromadia perennis*** (A.Gray) Greene: MEXICO, Baja California, Municipio Ensenada (Colonet Mesa), *Donald W. Kyhos s.n.* (DAV). ***Centromadia pungens*** subsp. ***pungens*** (Hook. & Arn.) Greene: USA, California, San Luis Obispo Co., *BGB 534* (DAV). ***Constancea nevinii*** (A.Gray) B.G.Baldwin: USA, California, Los Angeles Co. (Santa Catalina Island), *Steven A. Junak Sca-833* (JEPS). ***Deinandra fasciculata*** (DC.) Greene: USA, California, San Luis Obispo Co., *BGB 533* (DAV). ***Deinandra floribunda*** (A.Gray) Davidson & Moxley: USA, California, San Diego Co., *Fred Sproul s.n.*, 1988 (JEPS). ***Dubautia arborea*** (A.Gray) D.D.Keck: USA, Hawaii, Hawaii Island, *BGB 527b* (DAV). ***Dubautia herbstobatae*** G.D.Carr: USA, Hawaii, Oahu, *Gerald D. Carr 1244* (HAW). ***Dubautia knudsenii*** Hillebr. subsp. ***filiformis*** G.D.Carr: USA, Hawaii, Kauai, *Kenneth R. Wood 13268* (PTBG). ***Dubautia laevigata*** A.Gray: USA, Hawaii, Kauai, *BGB 777* (DUKE). ***Dubautia latifolia*** (A.Gray) A.Gray: USA, Hawaii, Kauai, *BGB 675* (DAV). ***Dubautia paleata*** A.Gray: USA, Hawaii, Kauai, *Gerald D. Carr 1375* (HAW). ***Dubautia plantaginea*** Gaudich. subsp. ***plantaginea***: USA, Hawaii, Maui (East Maui), *Hank Oppenheimer 11004* (UC). ***Dubautia scabra*** subsp. ***scabra*** (DC.) D.D.Keck: USA, Hawaii, Hawaii Island, *BGB 526* (DAV). ***Eatonella nivea*** A.Gray: USA, Nevada, Humboldt Co., *BGB 868* (UC). ***Eriophyllum lanatum*** subsp. ***obovatum*** (Pursh) J.Forbes: USA, California, San Bernardino Co., *Erin Shapiro s.n.*, 2008 (JEPS). ***Eriophyllum lanosum*** A.Gray: USA, California, San Bernardino Co., *BGB 888* (JEPS). ***Harmonia doris-nilesiae*** B.G.Baldwin: USA, California, Trinity Co., *Barbara Williams 518* (DAV). ***Harmonia guggolziorum*** B.G.Baldwin: USA, California, Mendocino Co., *Jack & Betty Guggolz 1635* (JEPS). ***Harmonia hallii*** B.G.Baldwin: USA, California, Colusa Co., *BGB 599* (DAV). ***Harmonia nutans*** (Greene) B.G.Baldwin: USA, California, Napa Co., *BGB 652* (DAV). ***Harmonia stebbinsii*** B.G.Baldwin: USA, California, Trinity Co., *BGB 611* (DAV). ***Hemizonia congesta*** DC.: USA, California, San Luis Obispo Co., *BGB 532* (DAV). ***Hemizonia congesta*** subsp. ***congesta*** DC.: USA, California, Sonoma Co., *Greg English-Loeb EL-2-Hc* (JEPS). ***Holocarpha macradenia*** (DC.) E.Greene: USA, California, Monterey Co., *BGB 724* (DAV). ***Holocarpha obconica*** (Nutt.) E.Greene: USA, California, Alameda Co., *BGB 1401* (JEPS). ***Holozonia filipes*** (Hook. & Arn.) Greene: USA, California, El Dorado Co., *BGB 513* (DAV). ***Hulsea algida*** A.Gray: USA, California, Alpine Co., *BGB 678* (DAV). ***Hulsea californica*** Torr. & A.Gray: USA, California, San Diego Co., *BGB s.n.*, 1992 (JEPS). ***Jensia rammii*** (Greene) B.G.Baldwin: USA, California, El Dorado Co., *BGB 494* (DAV). ***Jensia yosemitana*** (A.Gray) B.G.Baldwin: USA, California, Tuolumne Co., *BGB 603* (DAV). ***Kyhosia bolanderi*** (A.Gray) B.G.Baldwin: USA, California, El Dorado Co., *BGB 509b* (DAV). ***Lagophylla minor*** (A.Gray) J.T.Howell: USA, California, Napa Co., *BGB 600* (DAV). ***Lagophylla ramosissima*** Nutt.: USA, California, Solano Co., *BGB 936* (JEPS). ***Lasthenia californica*** DC.subsp. ***californica***: USA, California, Mariposa Co., *BGB 830* (JEPS). ***Lasthenia glaberrima*** DC.: USA, California, Solano Co., *BGB 884* (JEPS). ***Layia gaillardioides*** (Hook. & Arn.) DC.: USA, California, Marin Co., *BGB 1072* (JEPS). ***Layia heterotricha*** (DC.) Hook. & Arn.: USA, California, Santa Barbara Co., *BGB 738* (DAV). ***Madia citriodora*** A.Gray: USA, California, Butte Co., *BGB 715* (DAV). ***Madia elegans*** D.Don: USA, California, Kern Co., *BGB 695* (DAV). ***Madia radiata*** Kellogg: USA, California, San Luis Obispo Co., *BGB 693* (DAV). ***Madia subspicata*** D.D.Keck: USA, California, Butte Co., *BGB 713* (DAV). ***Monolopia gracilens*** A.Gray: USA, California, Contra Costa Co., *BGB 1069* (JEPS). ***Monolopia major*** DC.: USA, California, Colusa Co., *BGB 1011* (JEPS). ***Osmadenia tenella*** Nutt.: USA, California, San Diego Co., *Gerald D. Carr 1365* (DAV). ***Pseudobahia bahiifolia*** (Benth.) Rydb.: USA, California, Madera Co., *BGB 945* (JEPS). ***Pseudobahia peirsonii*** (D.D.Keck) D.D.Keck: USA, California, Kern Co., *BGB 913* (JEPS). ***Raillardella argentea*** (A.Gray) A.Gray: USA, California, Alpine Co., *BGB 512c* (DAV). ***Raillardella pringlei*** A.Gray: USA, California, Siskiyou Co., *BGB 608b* (DAV). ***Syntrichopappus fremontii*** A.Gray: USA, California, Kern Co., *BGB 1614* (JEPS). ***Syntrichopappus lemmonii*** A.Gray: USA, California, Kern Co., *BGB 1611a* (JEPS). ***Venegasia carpesioides*** DC.: USA, California, San Luis Obispo Co., *BGB 893* (JEPS). ***Wilkesia gymnoxiphium*** A.Gray: USA, Hawaii, Kauai, *Winona Char 76.022* (HAW). ***Wilkesia hobdyi*** H.St.John: USA, Hawaii, Kauai, *Gerald D. Carr 1150* (HAW).

## Notes

https://doi.org/10.5061/dryad.cjsxksnnr

### Competing Interest Statement

The authors have declared no competing interest.

